# Cold storage reveals distinct metabolic perturbations in processing and non-processing cultivars of potato

**DOI:** 10.1101/661611

**Authors:** Sagar S Datir, Saleem Yousf, Shilpy Sharma, Mohit Kochle, Ameeta Ravikumar, Jeetender Chugh

## Abstract

Cold-induced sweetening (CIS) causes a great loss to the potato (*Solanum tuberosum* L.) processing industry wherein selection of potato genotypes using biochemical information through marker-trait associations has found to be advantageous. In the present study, we have performed nuclear magnetic resonance (NMR) spectroscopy-based metabolite profiling on tubers from five potato cultivars (Atlantic, Frito Lay-1533, Kufri Jyoti, Kufri Pukhraj, and PU1) differing in their CIS ability and processing characteristics at harvest and after one month of cold storage at 4°C. A total of 39 water-soluble metabolites were detected using ^1^H NMR. Multivariate statistical analysis indicated significant differences in metabolite profiles between processing and non-processing potato cultivars. Further analysis revealed distinct metabolite perturbations as induced by cold storage in both types of cultivars wherein significantly affected metabolites were categorized mainly as sugars, sugar alcohols, amino acids, and organic acids. Significant metabolic perturbations were used to carry out metabolic pathway analysis that in turn tracked 130 genes encoding enzymes (involved directly and/or indirectly) involved in CIS pathway using potato genome sequence survey data. Based on the metabolite perturbations, the possible relevant metabolite biomarkers, significantly affected metabolic pathways, and key candidate genes responsible for the observed metabolite variation were identified. Overall, studies provided new insights in further manipulation of specific metabolites playing a crucial role in determining the cold-induced ability and processing quality of potato cultivars for improved quality traits.

**Highlight:** Metabolomic profiling using 1D ^1^H-NMR and bioinformatics analysis of potato cultivars for the identification of metabolites and genes controlling biochemical pathways in cold-stored potato tubers

## Introduction

Potato (*Solanum tuberosum* L.) – an important staple non-grain vegetable food crop – is used globally for both processing and table purposes. Cold storage of potato tubers after harvesting is mandatory to reduce sprouting, prevent diseases, avoid losses due to shrinkage, extend post-harvest shelf life, and to ensure year-round supply of quality tubers for consumption (Bianchi *et al.*, 2014; Hou *et al.*, 2017). During cold storage the potato tubers exhibit the phenomenon of cold-induced sweetening (CIS), wherein rapid degradation of starch and sucrose hydrolysis has been associated with accumulation of reducing sugars (RS) such as glucose and fructose (Burton, 1969; Dale and Bradshaw, 2003). During the frying process, these RS react with free amino acids in a Maillard reaction to generate dark-pigmented products that are bitter and unsightly to consumers. In addition to this, one of the products of the Maillard reaction is acrylamide – a potent neurotoxin and carcinogen (Menéndez *et al.*, 2002; Mottram *et al.*, 2002; Hajirezaei *et al.*, 2003). Hence, CIS is considered as one of the critical parameters in potato production as well as in processing; and therefore identification and development of potato tubers resistant to CIS has become a priority in a number of potato breeding programs (Xiong *et al.*, 2002; Hamernik *et al.*, 2009; Colman *et al.*, 2017). It is necessary to identify and develop potato cultivars with CIS resistance along with good processing quality attributes to meet the challenges of a frequently-changing market, production circumstances, and improving their economic condition (Kaur and Aggarwal, 2014). In this regard, the metabolic stability of potato tubers during the cold storage period has been identified as one of the prime traits to be investigated for breeding programs worldwide (Sowokinos, 2001; Ali and Jansky, 2015), wherein selection of potato genotypes at early generations using biochemical information through marker-trait associations has been found to be advantageous (Slater *et al.*, 2014; Gupta, 2017).

The potato processing industry is becoming an emerging sector in India and therefore, the demand for processed potato products such as chips, French fries, flakes, etc. is increasing continuously (Rana and Pandey, 2007). Ideally, potato cultivars suitable for processing should possess high specific gravity and dry matter (DM) content along with low RS content (Kaur *et al.*, 2012; Kaur and Aggarwal, 2014). In this regard, commercially grown processing (Atlantic and Frito Lay 1533) and popular Indian non-processing (Kufri Jyoti and Kufri Pukhraj) potato cultivars (Kaur and Aggarwal, 2014) along with one locally grown cultivar (PU1) were used as model-systems to identify bio-markers for CIS. While, Atlantic and Frito Lay-1533 have been rated as the best varieties for processing purpose with good storability, Indian potato cultivars Kufri Pukhraj and Kufri Jyoti are used for table purpose due to their medium and average/poor storability, but have been found to be inferior for processing purposes due to high RS and low DM content (Kaur and Aggarwal, 2014; Aggarwal *et al.*, 2017; Kaur and Khurana, 2017; Raigond *et al.*, 2018).

We carried out nuclear magnetic resonance (NMR)-based untargeted metabolic profiling of five potato cultivars differing in their CIS abilities from freshly harvested potatoes and after one month of cold storage (4°C). The key objective in this study was to examine the differences in metabolic profiles of these cultivars (between processing, non-processing, and local) at harvest and after cold storage to further advance the knowledge of biochemical mechanisms underpinning the CIS trait. The study also aimed to identify known biochemical pathways and to reveal underlying genes that control metabolite accumulation after cold storage of potato tubers. Finally, we targeted to identify key metabolite biomarkers and candidate genes (based on the metabolomics data and pathway analysis) that can potentially be used in breeding programs for the development of new cultivars for CIS resistance and improved processing attributes thereby enhancing the potato tuber quality.

## Materials and Methods

### Plant Material

Two potato cultivars Atlantic and Frito Lay-1533 (FL-1533) (Pepsi Foods Pvt. Ltd. Channo, Sangrur) suitable for processing purpose and two Indian cultivars Kufri Jyoti and Kufri Pukhraj (Central Potato Research Institute, Shimla) with non-processing characteristics differing in their cold storage ability (Marwaha *et al.*, 2005; Kaur and Aggarwal, 2014; Sharma *et al.*, 2012) used in the present study were obtained from BT Company and Jai Kisan Farm Products and Cold Chains Pvt. Ltd, India, Pune. One locally grown potato cultivar (PU1) possessed high RS and poor storability (Datir *et al.*, 2019) was also included in the study (Supplementary Table S1).

### Potato Plantation and Harvesting

Tubers of 5 cultivars, namely Atlantic, FL-1533, Kufri Pukhraj, Kufri Jyoti, and PU1, were planted in triplicates in separate PB 5 Polythene bags containing (potting mix: 60% shredded pine bark, 20% crusher dust, cow dung, 20% soil supplemented with sand and slow release fertiliser) on 25^th^ June 2018, at the Department of Biotechnology, Savitribai Phule Pune University, Pune, India. Tubers from 15 bags were harvested in the second week of October 2018. Six tubers of each cultivar (two tubers from each triplicate) were transferred to individual paper bags and divided into two groups consisting of three tubers each (one from each triplicate) for sampling at two treatments, (a) fresh harvest (FH), and (b) after one month of cold storage at 4°C (CS). While the first group of three tubers (one from each triplicate) from each of the 5 cultivars – making a total of 15 tubers for treatment (a) – were immediately processed for metabolite extraction, a second group of three tubers (one from each triplicate) from each of the 5 cultivars – making a total of 15 tubers – were stored at 4°C for one month (treatment b). All the tubers were subjected to freeze-drying (Operon, FDB-5503, Korea) for one week before using for metabolite extraction.

### Metabolite extraction

The freeze-dried potato samples (from treatments a and b of five cultivars) were ground to a fine powder and were used for metabolite extraction. Briefly, approximately 200 mg of freeze-dried potato powder was re-suspended in 200 µl Phosphate Buffer Saline (PBS) in 1.5 ml tubes and vortexed for five minutes. To each tube, 400 µl ice cold methanol (Sigma, HPLC grade) was added, followed by vortexing for another 5 min. Samples were then stored at −20°C for 12 h. Post-incubation, the samples were centrifuged at 16,000 *g* (Eppendorf centrifuge 5415C, Hamburg, Germany) for 20 min at 4°C. The supernatants were transferred to fresh 1.5 ml Eppendorf tubes and were subjected for lyophilization (Operon, FDB-5503, Korea). There were 3 replicates for each of the 5 cultivars processed in two treatments making a total of 30 distinct samples. The lyophilized extracts of all the samples were reconstituted into 580 µl 100% NMR buffer (20 mM sodium phosphate, pH 7.4 in D_2_O containing 0.4 mM DSS (2,2-dimethyl-2-silapentane-5-sulfonic acid). For making buffer containing a known concentration of DSS, 17.46 ± 0.01 mg of DSS was weighed (Mol wt. 218.32 g/mol) and dissolved in 2000 µl ± 2 µl of phosphate buffer. This stock solution was then diluted to 100 fold resulting in a final buffer solution containing 87.30 ± 0.16 mg/L of DSS in solution, which corresponds to 399.9 ± 0.7 µM of DSS in the buffer. The samples were vortexed for 2 min at room temperature and centrifuged at 4000 *g* for 2 min. The supernatants were transferred to respective 5 mm NMR tubes for NMR data measurements.

### NMR Spectroscopy

All the NMR data was measured on a Bruker AVANCE III HD Ascend NMR spectrometer operating at 14.1 Tesla. This spectrometer has been equipped with pulsed-field gradients in x, y, and z directions (operating at 54 Gauss/cm), and Bruker high-performance shim system with 36 orthogonal shim gradients and integrated real-time shim gradient for 3-axis shimming. A cryogenically cooled quad-channel (^1^H/^13^C/^15^N/^31^P-^2^H) probe was used to pump radio frequencies and detection. All the NMR data was measured at 298 K controlled by the Bruker VT unit. Water-suppression pulse sequence from Bruker library (noesygppr1d) was used to measure all the ^1^H-NMR data, where water suppression was achieved by pre-saturating water using continuous wave irradiation at 5.56E-05 W during the inter-scan relaxation delay of 5 s, and employing spoiler gradients (Smoothed square shape SMSQ10.100, where G_1_ was with 50% power and G_2_ was with −10% power for 1 ms duration each). The data acquisition period of 6.95 s (including inter-scan delay) was used, giving a spectral width of 7200 Hz resolved in 32k data points. Sixty-four scans were used to average the signal recorded on each sample. ^1^H 90° pulse-width, receiver gain, and water-suppression parameters were kept invariant from sample to sample to rule out intensity variations while recording data on different samples. To help with assignment of metabolites, ^1^H-^1^H total correlation spectroscopy (TOCSY) experiment (using mlevesgppg pulse sequence from Bruker library) was measured with a 6000 Hz of spectral width resolved in 2048 × 1024 data points with 40 transients per increment. A Hartman-Hahn mixing time of 80 ms was employed for the TOCSY spin-lock using composite blocks of 90°-180°-90° pulses with 90° pulse width of 25 μs at 2.29 W of power. TOCSY data was recorded in States-TPPI mode and Smoothed square shaped (SMSQ10.100) gradients were used with 31% power (after the spin-lock period) and 11% power (before refocusing) for a duration of 1 ms.

### Metabolite Identification and Quantification

All of the NMR data were processed using Topspin (v3.5) software (www.bruker.com/bruker/topspin). ^1^H NMR raw data was multiplied with exponential function and zero-filled to 64K data points prior to Fourier transformation. All the spectra are manually phased and the baseline is corrected before subjecting to further analysis. ^1^H chemical shift was directly referenced to DSS resonance (δ=0 ppm at 25 °C). ^1^H-^1^H TOCSY was processed with a pure cosine function (SINE with SSB = 2) and zero-filled to 2048 and 1024 data points in F1 and F2 dimensions prior to subjecting the data to Fourier transformation. Multiple peak parameters including, chemical shift values, J-coupling values, line shape, and multiplicity information, in combination with BMRB and HMDB data bases were used to assign the peaks to respective metabolites. Chenomx NMR suite 8.1 software was used to carry out the ^1^H resonance assignment with a chemical shift tolerance of 0.05 ppm when comparing the data with BMRB/HMDB. Resonance assignment of metabolites was confirmed using ^1^H-^1^H TOCSY (Supplementary Fig. S1) cross peak pattern of individual metabolites containing coupled ^1^H spin systems via a semi-automated software, MetaboMiner. A sets of five resonances remained unassigned and have been duly marked as U1-U5 (Supplementary Fig. S2 and Supplementary Table S2).

After identification of metabolites, respective peaks were manually picked, integrated using Topspin v3.5, and converted to absolute concentrations of individual metabolites using Chenomx NMR suite 8.1 by comparing with the peak integrals from an external reference compound DSS of known concentration (400 µM). The absolute concentrations obtained above were then normalized using the dry weight obtained from the tuber mass used for metabolite extraction. The data matrix file was created using concentrations of metabolites as obtained above from 30 distinct samples. The lower limit of quantification achieved using above-mentioned NMR parameters was 0.25 μM for the methyl peak of DSS at a s/n ratio of 10.

### Metabolic pathway analysis, Blast similarity searching, gene identification, notation and location on potato chromosomes

Metabolic pathway analysis depicting significantly affected metabolites in cold-stored potato tubers was performed by comparing the primary metabolites based on KEGG and the reference pathway (Sowokinos, 2001; Malone *et al.*, 2006) using MetaboAnylst web tool (https://www.metaboanalyst.ca/). BLAST similarity searching, gene identification and location in the potato genome annotated to encode enzymes of biochemical pathways was retrieved from Potato Genome Sequencing Consortium (PGSC) (http://solanaceae.plantbiology.msu.edu/pgsc_download.shtml), National Center for Biotechnology Information (NCBI) database (http://www.ncbi.nlm.nih.gov/), Sol Genomics Network (https://solgenomics.net/), Phytozome version 12.1 (https://phytozome.jgi.doe.gov/pz/portal.html) and KEGG (https://www.genome.jp/kegg/) using key word searches. The gene IDs have been taken from PGSC. In the event where gene sequences were not identified from PGSC, NCBI IDs have been provided.

### Statistical analysis

Due to high dimensionality and large complexity (5 cultivars in triplicates in 2 processing conditions with each NMR sample having ~1000 ^1^H signals) of the metabolomics data, multivariate statistical analysis was performed. To predict the differences in nature and concentrations of metabolite in various cultivars in triplicates with treatments a and b, principal component analysis (PCA) was carried out using normalized metabolite concentration as input in the MetaboAnalyst web tool (www.metaboanalyst.ca). The input data table was normalized using the Pareto-scaling approach available in the MetaboAnalyst. Correlation between first two principal components was drawn as the scores plot for all the samples and clusters of normal distribution were marked using ellipses showing 95% confidence limits for each group in PCA analysis. Next, pair-wise analysis of all five cultivars in FH and CS treatments was achieved using Volcano plot analysis, where metabolites were selected based on dual criteria, 1) the significance (false discovery rate (FDR) corrected p-value < 0.05), and fold-change in concentration (cut-off for fold change was set to 1.5 fold increase or decrease). In addition to this, the VIP score plot obtained by PCA identified the key metabolites responsible for the clustering of various groups. Metabolites with a VIP score of ≥1.0 are generally considered to be statistically significant (Ma *et al.*, 2016; Wu *et al.*, 2018). A union set of significant metabolites (those identified from volcano plot analysis, and from VIP score following the above-mentioned criteria) were taken for generating Box and Whisker plots to highlight the variation of a particular metabolite across replicates, different cultivars, and in different treatment conditions. Metabolites, e.g. ascorbate, having low signal-to-noise (s/n < 15) in NMR spectra, although identified with confidence, were not included in box and whisker plot analysis as they might be prone to over- or under-estimation of concentrations. Further, correlation plots were drawn to identify all the correlated metabolites in FH and CS treatments for all five cultivars. The significantly affected pathways were then identified using significantly perturbed metabolites as input in MetaboAnalyst tool and KEGG pathway database (www.genome.jp/kegg/pathway.html).

## Results and discussion

### Global profiling of metabolites in different potato cultivars – Processing versus non-processing cultivars

Global profiling of metabolites obtained from the methanolic extracts of the FH and CS tubers obtained from five different potato cultivars, differing in their cold storage behaviour and processing characteristics (Supplementary Table S1), identified a total of 39 abundant water-soluble metabolites using 1D ^1^H-NMR (Supplementary Fig. S2); and confirmed using 2D ^1^H-^1^H TOCSY (Supplementary Fig. S1) and BMRB database. Identified metabolites have been marked on 1D ^1^H-NMR (Supplementary Fig. S2), listed in supplementary table S2, and were quantified. Water-insoluble metabolites in the organic phase gave broad and overlapping signals in 1D ^1^H-NMR and thus were excluded from the analysis. A range of distinct metabolites was detected that could be characterized mainly as sugars, sugar alcohols, amino acids, and organic acids (Supplementary Table S2). The unsupervised PCA analysis showed a divergent separation on the scores plot of PC1 and PC2, accounting for a 55.4% and 22.2% variation in the metabolites extracted from the FH and CS treated tubers of the different cultivars used in the study (Fig. 1). Interestingly, the clustering of single points in the principal component space (marked by ellipses showing 95% confidence limits of a normal distribution) for metabolites from the FH tubers of Atlantic and FL-1533 (processing cultivars) clustered together while FH tubers from Kufri Jyoti and Kufri Pukhraj cultivars and the local PU1 were more similar to each other (Fig. 1). Further, the ellipses marking the principal component space for the metabolites of the cold storage tubers were also found to be different in processing, non-processing, and locally grown cultivars (Fig. 1). These differences in the metabolite content in the different cultivars at the two time-points could be attributed to the genetic make-up of each cultivar used in the present study. In fact, previous studies have also reported such variability in the metabolite content from different potato cultivars differing in their genetic background (Defernez *et al.*, 2004; Uri *et al.*, 2014).

**Fig. 1:**
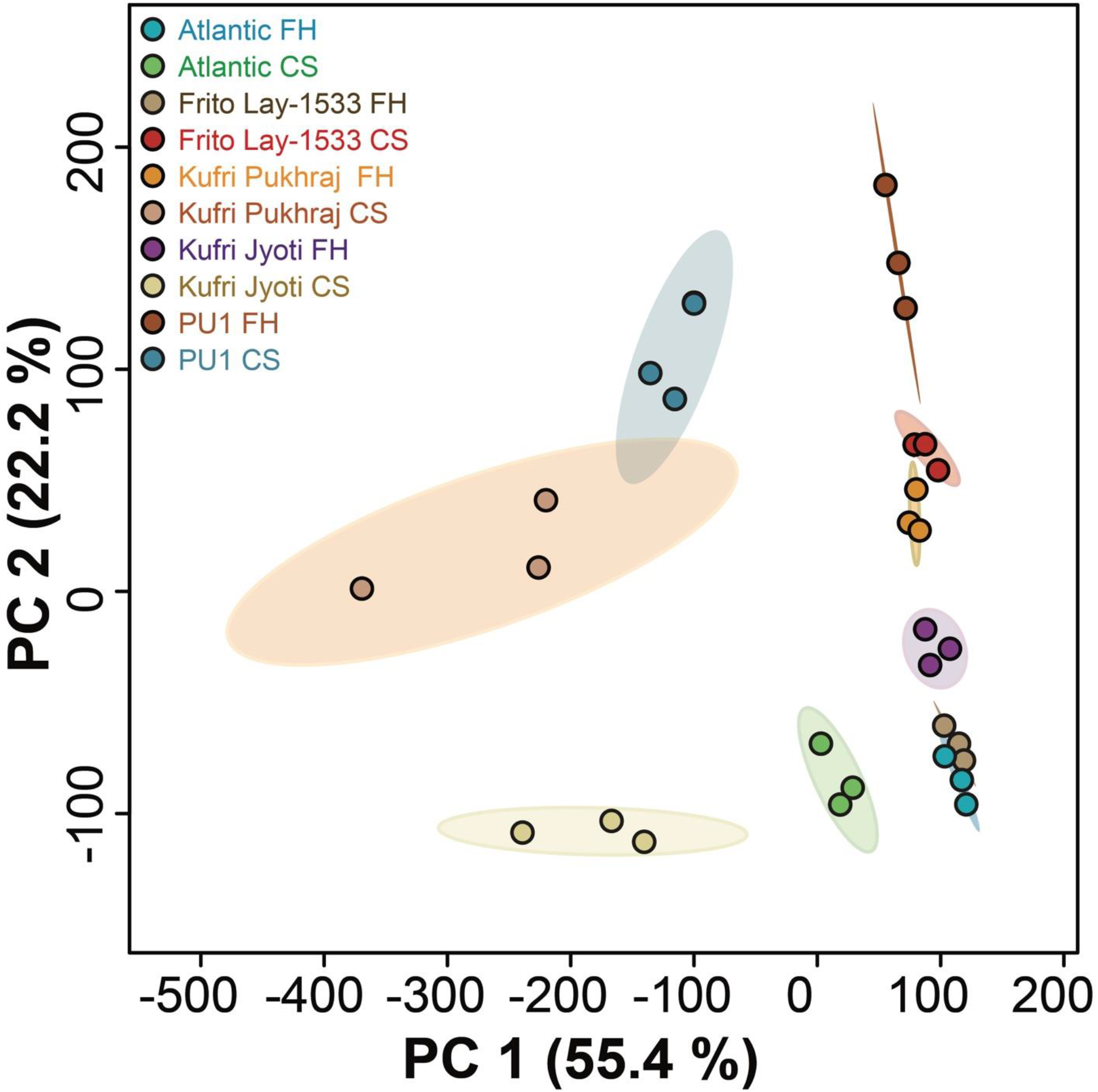
Scores plot as obtained by PCA utility of MetaboAnalyst software for the different potato cultivars (Atlantic, Frito Lay-1533, Kufri Pukhraj, Kufri Jyoti, and PU1) at fresh harvest (FH) and one month cold storage at 4°C (CS). Three replicates were used for each cultivar and at each condition (as described in Materials and Methods). Ellipses showing 95% confidence limits of a normal distribution for each group of the samples have been marked in respective colours for each cultivar. Color legends have been mentioned in the figure.

### Pair-wise analysis of metabolic changes upon cold storage in processing, non-processing, and local cultivars

The variations in metabolite profiles of potato cultivars differing in their genetic constitution offer a potential tool to develop CIS resistant potatoes with genotypes encoding improved processing characteristics. However, studies investigating the metabolic diversity from cold-stored potato tubers differing in their processing and non-processing characteristics have been limiting. In order to highlight the differences in the FH and the CS condition from the processing, the non-processing, and the local potato cultivars used in the study, pair-wise analysis was done (Fig. 2). In addition to this, volcano plot analysis (Fig. 3) and VIP score plot analysis (Fig. 4) was also performed under these conditions to identify the significantly affected metabolites in cold storage.

**Fig. 2:**
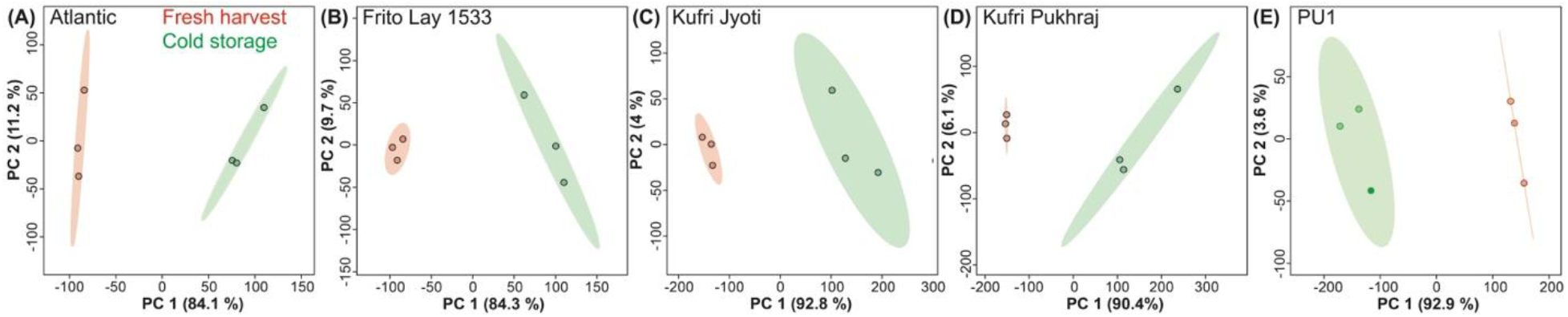
PCA score plots for pair-wise analysis of metabolites obtained from the different cultivars of potatoes at fresh harvest (Red) and cold storage at 4°C for 1 month (Green). A) Atlantic, B) Frito Lay-1533, C) Kufri Jyoti, D) Kufri Pukhraj, and E) PU1. Ellipses showing 95% confidence limits of a normal distribution for each group of the samples have been marked in respective colours (as mentioned above).

**Fig. 3:**
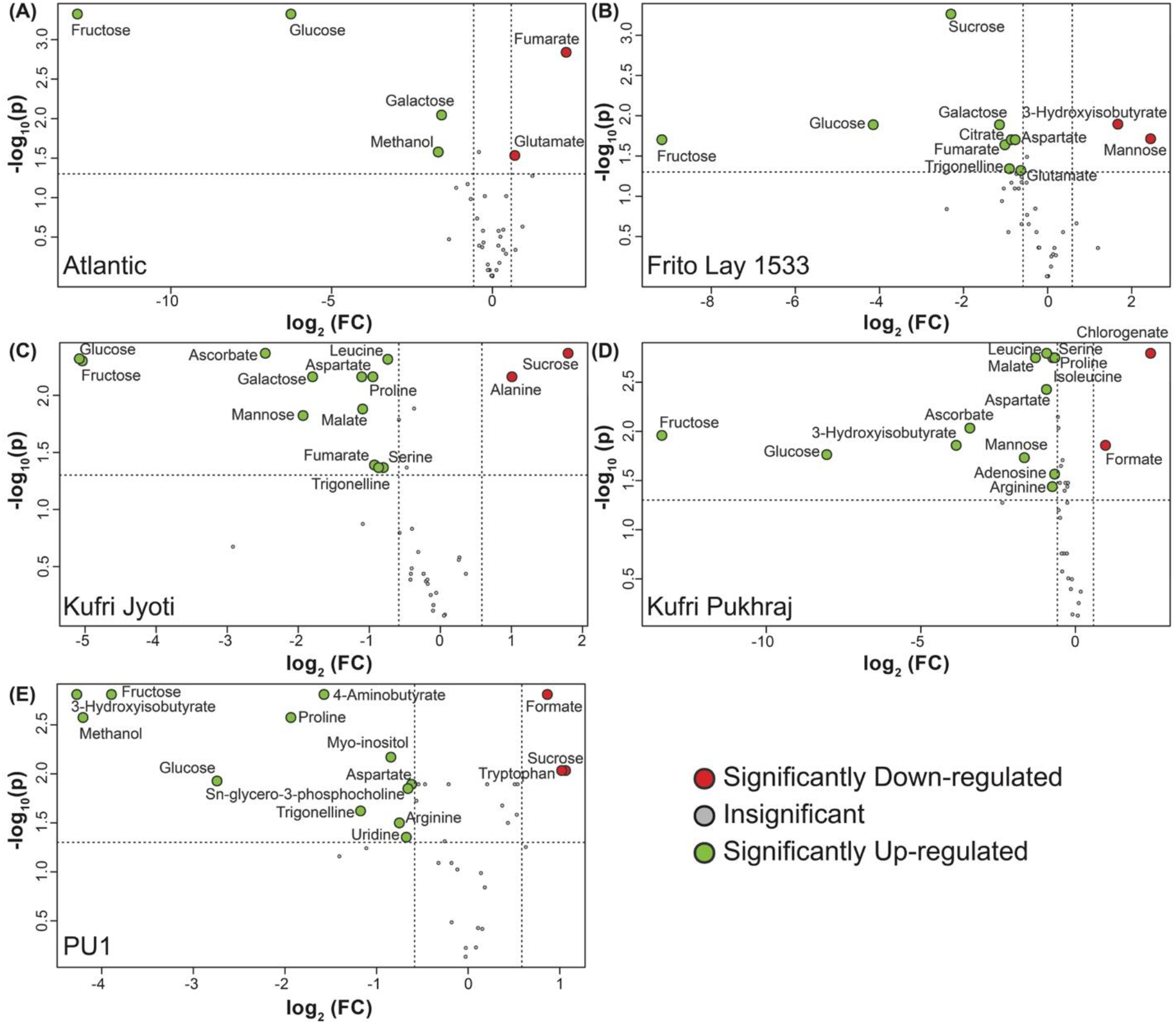
Volcano plots, where log_10_(FDR-corrected p-value) is plotted against log_2_(fold-change in concentration), depicting the changes in the metabolite concentration from freshly harvested potato tubers and tubers stored at 4 °C for one month. The different cultivars used for the study have been depicted as A) Atlantic, B) Frito Lay-1533, C) Kufri Jyoti, D) Kufri Pukhraj, and E) PU1. The significantly down-regulated metabolites upon cold storage have been marked in red and the ones up-regulated have been marked in green.

**Fig. 4:**
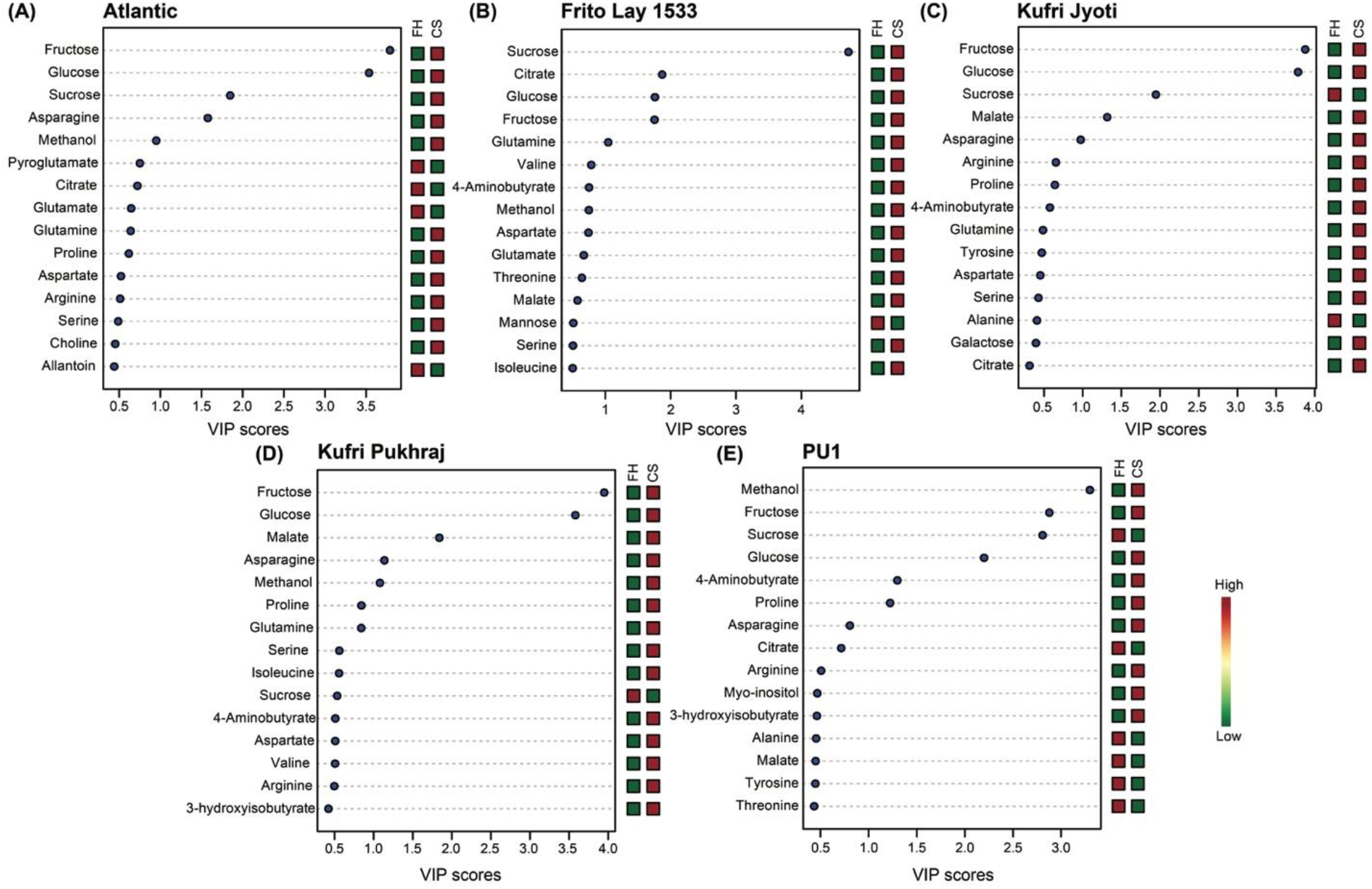
VIP scores obtained after pair-wise PCA analysis for A) Atlantic, B) Frito Lay-1533, C) Kufri Jyoti, D) Kufri Pukhraj, and E) PU1. A VIP score of ≥1.0 is considered significant.

### a. Metabolic changes in processing cultivars in CS conditions

The chemometric analysis was performed to assess the metabolic perturbations of potato tubers upon CS in all 5 cultivars. PCA analysis of metabolites obtained from the processing cultivars, Atlantic and FL-1533 showed 84.1% variation in PC1 and 11.2% variation in PC2 (Fig. 2A); and 84.3% variation in PC1 and 9.7% variation in PC2 (Fig. 2B), respectively. While in the Atlantic cultivar, fumarate and glutamate were found to be significantly downregulated upon CS when compared with FH; fructose, glucose, galactose, methanol, sucrose, and asparagine were significantly upregulated upon CS (Fig. 3A and Fig. 4A). Similarly, in the FL-1533 cultivar, CS treatment significantly increased the levels of fructose, glucose, sucrose, galactose, fumarate, trigonelline, citrate, aspartate, glutamate, and glutamine (Fig. 3B and Fig. 4B). On the other hand, the levels of mannose and 3-hydroxyisobutyrate were significantly reduced upon CS in this cultivar.

### b. Metabolic changes in non-processing and the local cultivars in cold storage conditions

PCA analysis of metabolites obtained from the non-processing cultivars, Kufri Jyoti and Kufri Pukhraj, showed 92.8% variation in PC1 and 4% variation in PC2 (Fig. 2C); and 90.4% variation in PC1 and 6.1% variation in PC2 (Fig. 2D), respectively upon CS. The levels of glucose, fructose, mannose, galactose, aspartate, malate, fumarate, leucine, proline, trigonelline, asparagine, and serine were increased in the Kufri Jyoti cultivar upon CS, while the levels of sucrose and alanine were reduced (Fig. 3C and Fig. 4C). Similarly, CS treatment of the Kufri Pukhraj cultivar was associated with significant increase in the levels of fructose, glucose, 3-hydroxyisobutyrate, mannose, malate, leucine, aspartate, serine, proline, isoleucine, adenosine, arginine, asparagine, and methanol on one hand; it significantly decreased the levels of chlorogenate and formate upon CS (Fig. 3D and Fig. 4D). In the local PU1 cultivar, levels of formate, tryptophan, and sucrose were significantly decreased, while 3-hydroxyisobutyrate, methanol, fructose, glucose, proline, 4-aminobutyrate, trigonelline, myo-inositol, arginine, aspartate, uridine, and sn-glycero-3-phosphocholine showed significant increase upon cold storage treatment (Fig. 3E and Fig. 4E).

Metabolomics approach has been previously used to assess the effect of storage conditions on a variety of potato cultivars. For example, metabolic profiles in different life cycle stages of potato tubers were characterized to link temporal changes in metabolites related to trait development (Shepherd *et al.*, 2010). In a recent study, comprehensive metabolomics and ionomics analysis on raw and cooked potato tubers of 60 unique genotypes were performed to understand the chemical variation and nutritional values in different varieties (Chaparro *et al.*, 2018). In another study, storage of commercial cultivars at 20-22 °C in the dark suggested a significant decrease in sucrose and fructose (Uri *et al.*, 2014). Here, we have reported that the storage of potato tubers at 4°C significantly increased the levels of sucrose, particularly in Atlantic and Frito Lay 1533, while it was significantly decreased in Kufri Jyoti and PU1, and remained invariant in Kufri Pukhraj (Fig. 5). On the other hand, we found that the increase in RS was more pronounced in the non-processing cultivars Kufri Pukhraj, Kufri Jyoti, and PU1 as compared to the processing cultivars, Atlantic and Frito Lay 1533 (Fig. 3, Fig. 4, and Fig. 5). These results are in agreement with other studies that observed an increase in RS after cold storage of potato tubers (Kaur and Aggarwal, 2014; Aggarwal *et al.*, 2017; Datir *et al.*, 2019), which has been attributed to the enhanced activity of the vacuolar invertase (Lin *et al.*, 2013). The effect of silencing of vacuolar invertase, which converts sucrose into glucose and fructose, on sugar metabolism pathways has previously been studied to find suitable targets for further genetic manipulation to improve the tuber quality (Wiberley-Bradford *et al.*, 2014). Brummell *et al.*, (2011) analysed the RS along with the expression of invertase and invertase inhibitors in cold-stored potato tubers obtained from cold-sweetening susceptible and cold-sweetening resistant cultivars. They demonstrated that the levels of RS decreased after one month of cold storage and this was accompanied by an increase in expression of the vacuolar invertase inhibitor mRNA accumulation in processing cultivars. Therefore, a relatively lower increase of RS after cold storage in Atlantic and Frito Lay 1533 cultivars (when compared with non-processing and local cultivar) used in this study could be attributed to increased levels of vacuolar invertase inhibitor. We recently studied the allelic variations in the vacuolar invertase inhibitor gene from Atlantic, Frito Lay 1533, Kufri Jyoti, Kufri Pukhraj, and PU1 cultivars and proposed that the SNPs in the vacuolar invertase inhibitor gene could be associated with the variation in RS levels in these cultivars (Datir *et al.*, 2019). However, these results need to be further validated using a qRT-PCR expression of vacuolar invertase inhibitor gene before and after cold storage in these cultivars.

**Fig. 5:**
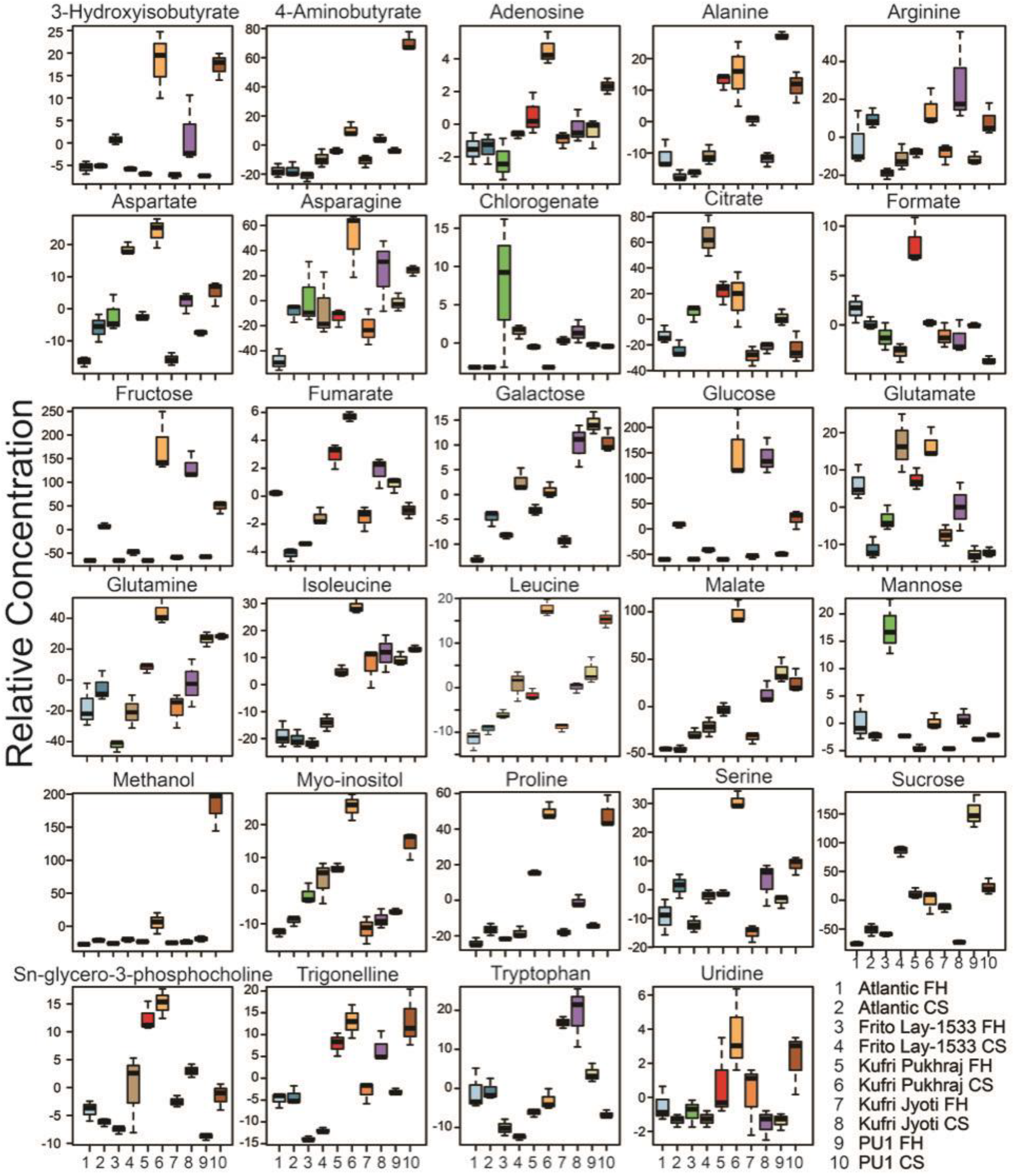
Box-Whisker plot for the significantly different metabolites (p-value < 0.05, and with VIP score ≥1.0) for the different potato cultivars. The significantly different metabolites obtained from ANOVA and post-hoc analysis were selected individually and the relative concentrations of each of these were plotted against the two time-points, i.e., fresh harvest and one month cold storage for the 5 cultivars used in the study. FH – fresh harvest and CS – cold storage at 4°C.

Although CS resulted in several significant metabolic perturbations, it is important to highlight the significance of some metabolites that can be related to the CIS status of the potato cultivars. For example, it is noteworthy that FL-1533 exhibited significantly higher citrate levels as compared to rest of the cultivars after CS (Fig. 5), which might be associated with CIS resistance along with chips with an acceptable colour. This is particularly important as citric acid is known as a popular anti-browning agent, mainly because it not only inhibits the polyphenol oxidase by reducing pH but also chelates copper at the enzyme-active site (McCord and Kilara 1983). Likewise, the changes in the levels of total amino acids, specifically the levels of asparagine, and the ratio of free asparagine to RS during cold storage were found to be significantly varied among different potato cultivars upon CS (Fig. 5). These factors, therefore, can further influence the processing quality of potato tubers. It is interesting to note that, among all the cultivars, PU1 cultivar in particular showed significantly higher levels of methanol after CS (Fig. 5). The amount of methanol released on saponification is the measure of the degree of pectin methylation and was found to be associated indirectly with the potato tuber texture properties (Ross *et al.*, 2010b). It can also be presumed that some of these significantly affected metabolites might have acted as osmolytes such as proline, trigonelline, 4-aminobutyrate (GABA), etc. (Evers *et al.*, 2010) (Fig. 5, Fig. 6) as an acclimation response under CS treatment. However, not much research has been focused on understanding the function of various metabolites in the CIS process of potato tubers. Therefore, these uniquely observed metabolite variations provide new insights into identifying and developing CIS resistant potato genotypes.

**Fig. 6:**
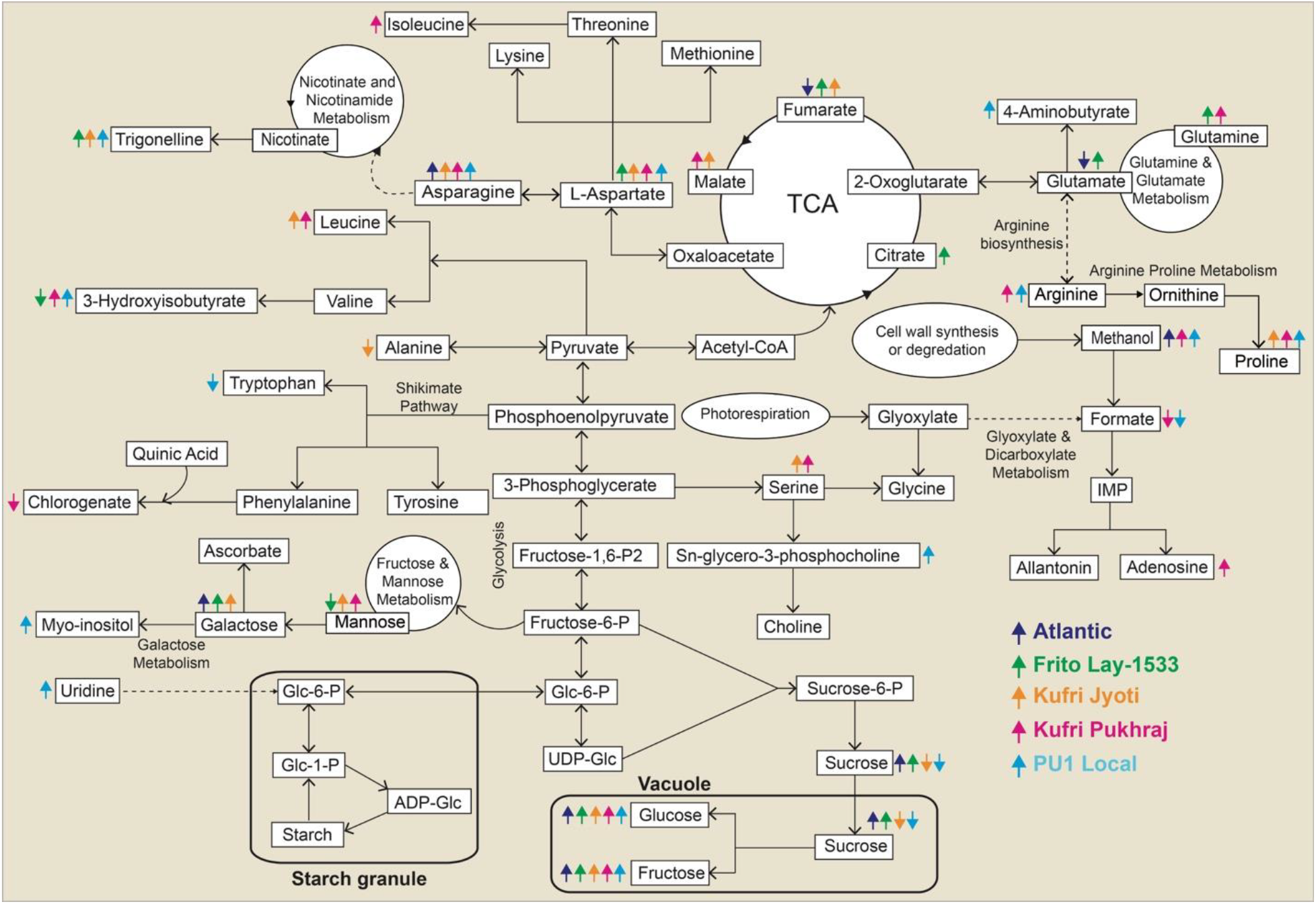
Pictorial representation of metabolic pathways affected during cold-induced sweetening in the different cultivars of potato. The significantly different metabolites under cold storage have been marked with arrows wherein an ↑ indicates upregulated metabolites and the ↑ arrow indicates down regulated metabolites. TCA – tricarboxylic acid.

### Metabolic correlation network analysis

Person’s correlation coefficient analysis was used to analyze the metabolite-metabolite correlation among identified metabolites in all five cultivars at both the time-points (Supplementary Fig. S3-S7). A total of 100, 84, and 45 significant correlations (p-value < 0.5) were obtained at FH among processing group (Atlantic and FL-1533), non-processing group (Kufri Pukhraj and Jyoti), and local (PU1) cultivars, respectively (upper-right half of the plot marked with white triangle in supplementary Fig. S3-S7). After CS, the number of significant correlations changed to 108, 105 and 31 (lower-left half of the plot marked with blue triangle in supplementary Fig. S3-S7) in the processing group, non-processing group, and local cultivar, respectively. The number of positive vs. negative correlations also varied depending on the variety (Supplementary Table S3). Remarkably, among all the metabolites, amino acids dominated the significant metabolite correlations. In general, the metabolite-metabolite correlations detected in the present work were highly dependent on the type of the cultivar considered; however, some particular behaviours of the metabolic network after CS are worth mentioning. For instance, a positive correlation of fructose with phenylalanine, valine, 4-aminobutyrate, glutamine, choline, and glutamate was evident in Atlantic (Supplementary Fig. S3) upon CS. In the case of FL-1533, pyroglutamate was found to be positively correlated with valine, alanine, arginine, and 4-aminobutyrate after CS (Supplementary Fig. S4). Whereas 4-amiobutyrate was positively correlated with trigonelline, sucrose, allantoin, and arginine, citrate and malate displayed significantly positive correlations with several other metabolites in Kufri Jyoti after CS (Supplementary Fig. S5) which were not evident in the other cultivars. However, several negative correlations were exclusively observed in PU1 cultivar after CS (Supplementary Fig. S7). No correlation between metabolites that are close in a metabolic pathway was observed after CS. For instance, glutamate and glutamine are metabolic neighbours in the glutamine synthase pathway and are found to be uncorrelated in the non-processing and local cultivars (Supplementary Fig. S5-S7) in FH as well as CS treatments, while are found to be correlated in processing cultivars (Supplementary Fig. S3 and S4). On the other hand, several other metabolite correlations were noted even if they are not metabolic neighbours. Significant correlations among various potato cultivars might help to predict the CIS status of the particular potato genotype based on FH and CS tuber profiling. However, the reason for these strong correlations remains unclear as no direct link has been reported so far and further investigation is needed. Metabolite correlations of potato groups differing in the genetic background have been previously reported (Dobson *et al.*, 2010; Chaparro *et al.*, 2018). Significant metabolite variation and metabolite-metabolite correlations were detected from a collection of 60 unique potato genotypes that span 5 different market classes such as russet, red, yellow, chip, and speciality (Chaparro *et al.*, 2018), where authors concluded that metabolite diversity and correlations data can support the potential to breed new cultivars for improved health traits.

### Metabolite biomarkers for the identification of CIS resistant and susceptible genotypes

CIS is a multigenic complex trait involving multiple intricate metabolic pathways which clearly indicates that it is unlikely to be controlled by a single metabolite; thus, multiple metabolites would come-up as plausible biomarkers for CIS in potatoes. Previous studies have suggested that various primary metabolites in potato tubers can be utilized as biomarkers in breeding programs for predicting agronomically important traits such as black spot bruising and chip quality (Steinfath *et al.*, 2010; Instroza-Blancheteau *et al.*, 2018). We would like to point out that in addition to the amount of RS, the total and individual amino acid content, the asparagine content, levels of organic acids, and other metabolites could be considered as important processing parameters. Breeders aim for the identification and development of processing potato cultivars with low free-asparagine and RS as desirable characteristics for processing purpose. In the current study, a unique metabolite combination was observed for the processing cultivar, FL-1533, which was represented by the lowest amount of RS and asparagine compared to rest of the cultivars CS (Fig. 3, Fig. 4, and Fig. 5). The levels of RS and asparagine have been used as markers for potato trait development (Shepherd *et al.*, 2010).

Amongst the different TCA cycle metabolites, the levels of citrate were found to be significantly higher in the processing cultivar, FL-1533, whereas Kufri Jyoti and Kufri Pukhraj showed significantly higher levels of malate after CS (Fig. 6). Citrate and malate are critical in determination of non-enzymic browning reactions, after cooking darkening, physiological age/ stages of development in the storage of potato tubers (Wichrowska *et al.*, 2009; Reust and Aerny, 1985). In addition, they indirectly influence the texture of cooked and fried potato products (Heisler *et al.*, 1964; Thomas *et al.*, 1979; Lynch and Kaldy, 1985). Hence, it is necessary to develop the indicators of tuber browning and physiological age mainly because both the growers and seed companies can optimize the storage conditions for individual cultivars. Moreover, such indicators will be extremely important in the determination of the suitability of potato tubers for culinary use and industrial processing (Reust and Aerny, 1985).

The texture of potato tubers is a key determinant of the quality of processing as well as cooked potato as has been shown to greatly influence the consumer’s preference (Shomer and Kaaber, 2006; Thybo *et al.*, 2006; McGregor, 2007). It is mainly determined by the breakdown of the cell wall middle lamella during cooking, and the correlation between pectin methylesterase activity and the degree of methylation of cell wall pectin (reviewed in Taylor *et al.*, 2007; Ross *et al.*, 2010b). The amount of methanol released on saponification is the measure of the degree of pectin methylation and is indirectly associated with the potato tuber texture properties (Ross *et al.*, 2010b). Significantly highest levels of methanol were exclusively recorded in the PU1 cultivar after CS (Fig. 5). Therefore, the amount of methanol present in potato tubers can be used as a potential marker for screening of potato cultivars for texture properties.

Importantly, several other metabolites such as fumarate, adenosine, sn-glycero-3-phosphocholine, 4-aminobutyrate, 3-hydroxyisobutyrate, trigonelline, and chlorogenate were significantly varied upon CS (Fig. 3, Fig. 4, and Fig. 6) indicating that these metabolites might have a role in the CIS process, as well as the determination of processing quality of these cultivars. In order to improve potato (*Solanum tuberosum* L.) genotypes through selection or breeding, it is helpful to determine the chemical composition of tubers (Pal *et al.*, 2008). Maintaining the quality of potato tubers during storage is a major challenge. Therefore, the information on the response of potato cultivars to cold storage and metabolite accumulation can be useful for the development of biomarkers predicting severity of CIS of different potato genotypes. Such biomarkers (supplementary table S4) can then be tested on a wide range of potato genotypes differing in CIS response and easily integrated into the existing potato storage management and breeding methods. Moreover, such predictive biomarkers can be used in selection for potato breeding and for tailoring storage conditions for each lot of harvested tubers (Neilson *et al.*, 2017). Furthermore, biomarkers can be utilized for the manipulation of a specific metabolite pathway for developing potato genotypes with improved processing characteristics.

### Metabolic Pathway Analysis

We performed the pathway analysis depicting significantly affected metabolites in cold-stored potato tubers by comparing the primary metabolites based on KEGG and the reference pathway (Fig. 6) (Sowokinos, 2001; Malone *et al.*, 2006). Cold temperature induces starch degradation in potato tubers to principal sugars including sucrose, glucose, and fructose, thereby leading to an imbalance between starch degradation and sucrose metabolism in tubers. So far, CIS studies have mainly concentrated on the activity of enzymes involved in the conversion of starch and sugars (Jansky and Fajardo, 2014). However, potato tubers displayed diverse biochemical mechanisms during CIS and the amount of sugar in potato tubers is influenced by several candidate genes operating in glycolysis, hexogenesis, and mitochondrial respiration (Sowokinos, 2001). The metabolic pathway analysis presented in this study suggests that several metabolites were affected during cold storage and mainly resulted from the alanine, aspartate, and glutamate metabolism; valine, leucine, and isoleucine biosynthesis; arginine and proline metabolism; glycine, serine, and threonine metabolism; the TCA cycle, fructose and mannose metabolism, galactose metabolism, nicotinate and nicotinamide metabolism, glycolysis; and sucrose metabolism along with several other metabolites (Fig. 6). Also, the levels of metabolites were found to be specifically different depending on potato cultivars (Fig. 3, Fig. 4, and Fig. 5) indicating that the specific metabolites might play a crucial role in determining the cold-induced ability of potato cultivars. Also, the molecular events controlling such metabolic perturbations in potato tubers after cold storage are still puzzling. Among various metabolic processes, carbohydrates, amino acids and organic acids were identified as the main players in the CIS process and were either decreased or increased under cold storage condition. In the amino acid metabolism pathways, the distinct significantly affected pathways include metabolism of 11 amino acids: isoleucine, glutamate, glutamine, leucine, alanine, arginine, proline, tryptophan, aspartate, asparagine and serine metabolism. In the TCA cycle, the levels of citrate, malate, and fumarate were significantly affected by CS. Particularly, citrate and fumarate synthesis was up-regulated in FL-1533 cultivar (Fig. 6). Several other metabolites such as 3-hydroxyisobutyrate, trigonelline, galactose, mannose, etc. were either up-regulated or down-regulated in response to cold storage (Fig. 6). On the other hand, methanol production was significantly enhanced in Atlantic, Kufri Pukhraj, and PU1 cultivars although the extent of this increase was significantly higher in PU1 (Fig. 6). The GABA shunt pathway was significantly enhanced as seen by the increased levels of 4-aminobutyarte in PU1 upon cold storage, whereas it was not significantly affected in any other potato cultivar. The convergence and divergence of various pathways involved in CIS revealed a complex metabolic network. However, the roles of these metabolites and their accumulation pattern in response to cold storage in different potato cultivars remains to be further investigated. A possible approach to achieve this goal is to identify the genes which are putatively involved in the formation of enzymes involved in the biosynthesis of these metabolites.

### Putative genes controlling CIS process of potato tubers

Taking leads from the identified metabolites and the major metabolic pathways affected during CIS, key-word searches for the gene name were conducted on multiple databases including PGSC, NCBI, Sol Genome Network, and Phytozome to identify candidate genes likely to be involved in the observed metabolic variation (Table 1). Based on the metabolic pathway analysis, a total of 29 significantly affected metabolites (Fig. 6) encompassing 130 genes that are likely to participate in CIS mechanism were identified (Table 1). Although, candidate genes involved in starch and sugar metabolism linked to the quantitative trait loci (QTL) for sugar and starch contents have been reported earlier (Chen *et al.*, 2001), information about several other enzymes controlling various metabolites in CIS is still lagging, which lays an obstacle for metabolic engineering of potato. Fischer *et al.*, (2013) reported that besides starch-sugar interconversion and membrane composition, the adaptation of tubers to cold storage might include other pathways. In the current study, in addition to sucrose metabolism, majority of the enzymes involved in controlling amino acid metabolism, and organic acids metabolism were identified (Table 1). In several cases, more than one isoform was identified indicating most of the enzymes were encoded by multigene families. Interestingly, tryptophan synthase, sucrose phosphate phosphatase, sucrose synthase, malate dehydrogenase, glutamine synthetase related to tryptophan, sucrose biosynthesis, malate, and glutamine metabolism represented by a larger multigene families consisting of 5 copies of genes located on various chromosomes. Also, several genes annotated with different loci were observed (Table 1). Metabolic perturbations in response to cold storage (Fig. 3, Fig. 4, and Fig. 5) could be attributed to either the natural allelic variations of genes or changes in transcript levels. Natural variation in candidate genes such as invertase and invertase inhibitors revealed that the genetic polymorphism raises the possibility that SNPs in alleles of these genes may contribute to the phenotypic variation in response to CIS among the potato genotypes (Menéndez *et al.*, 2002; Baldwin *et al.*, 2011; Datir *et al.*, 2012; Datir *et al.*, 2019).

**Table 1:**
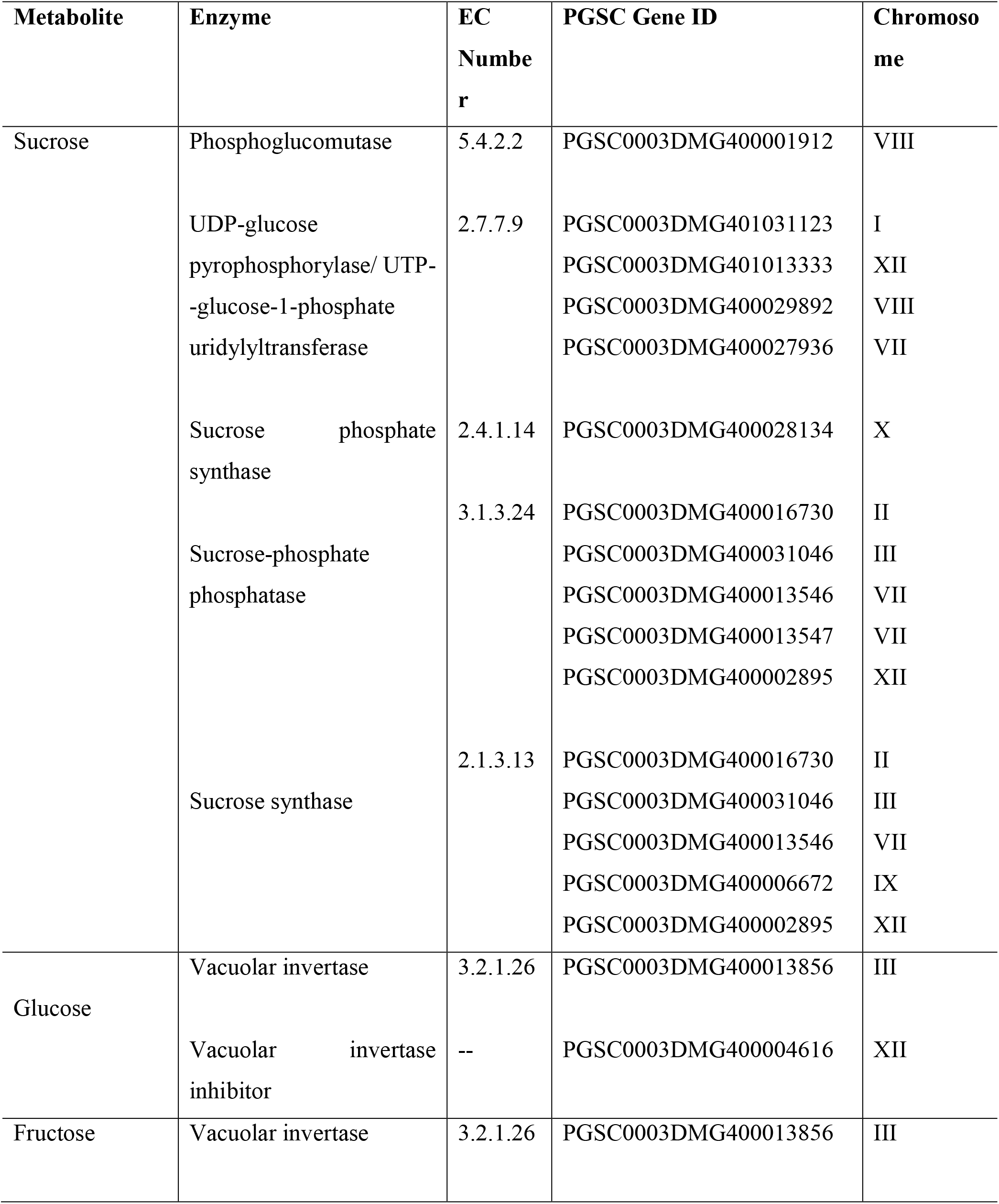
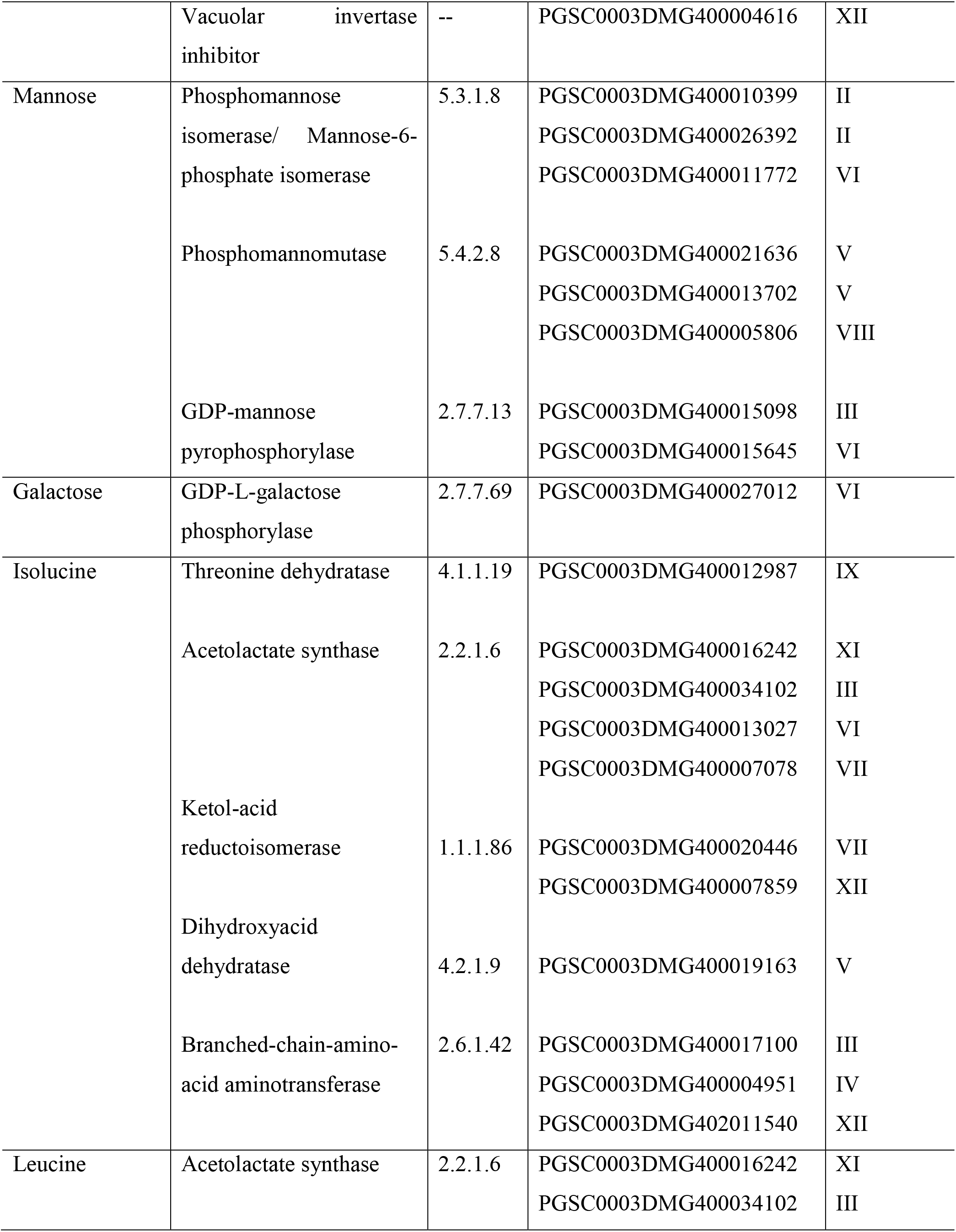
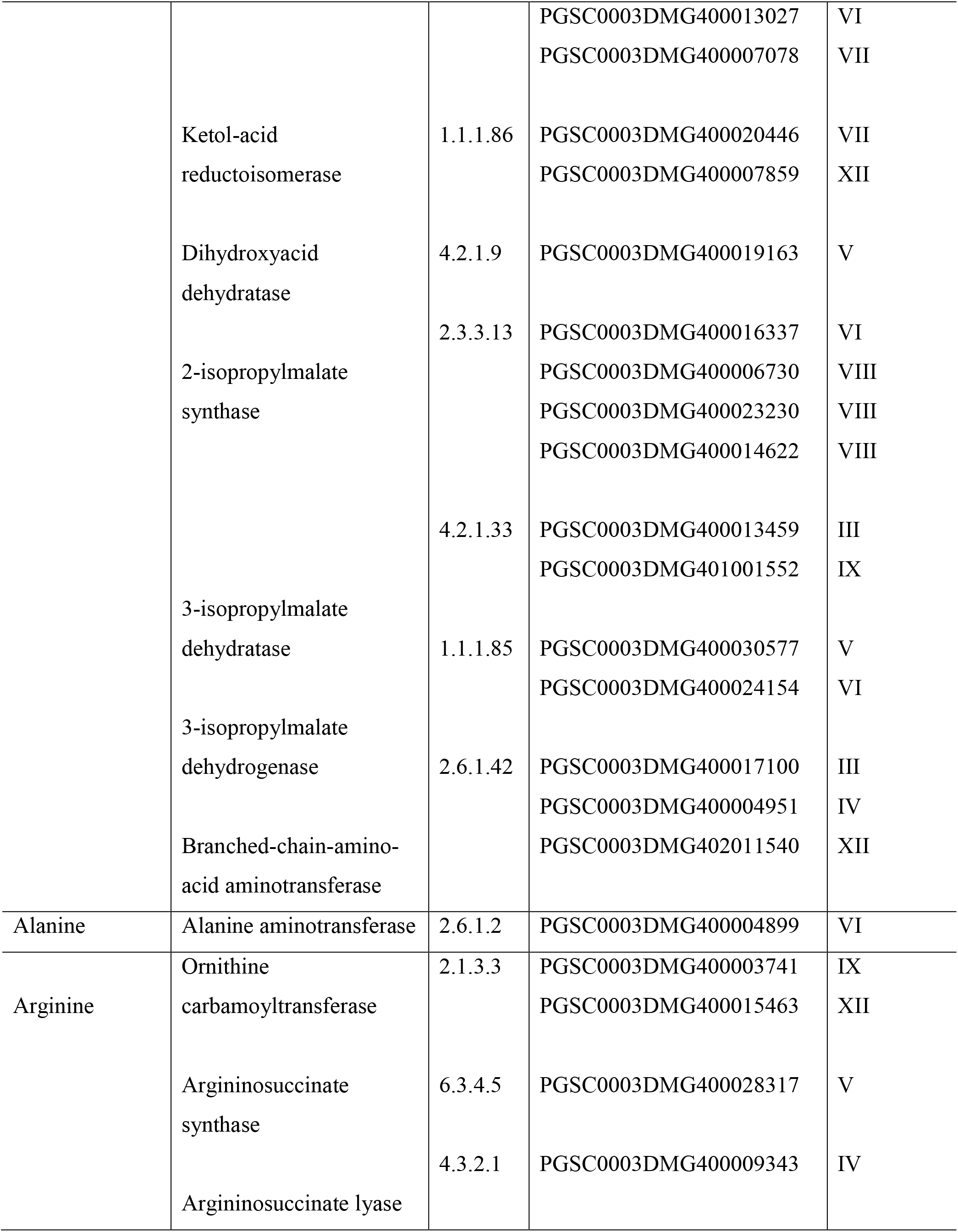
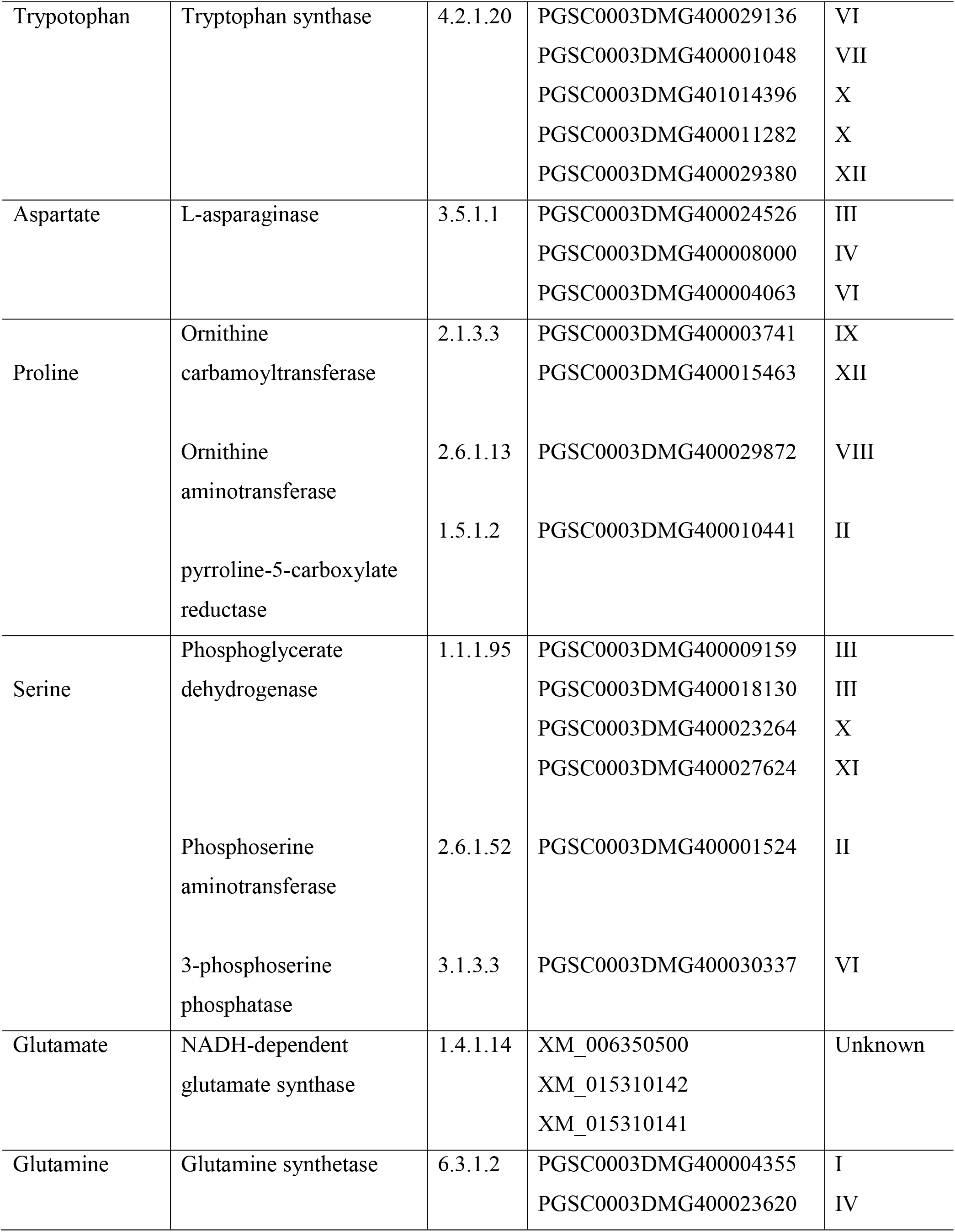
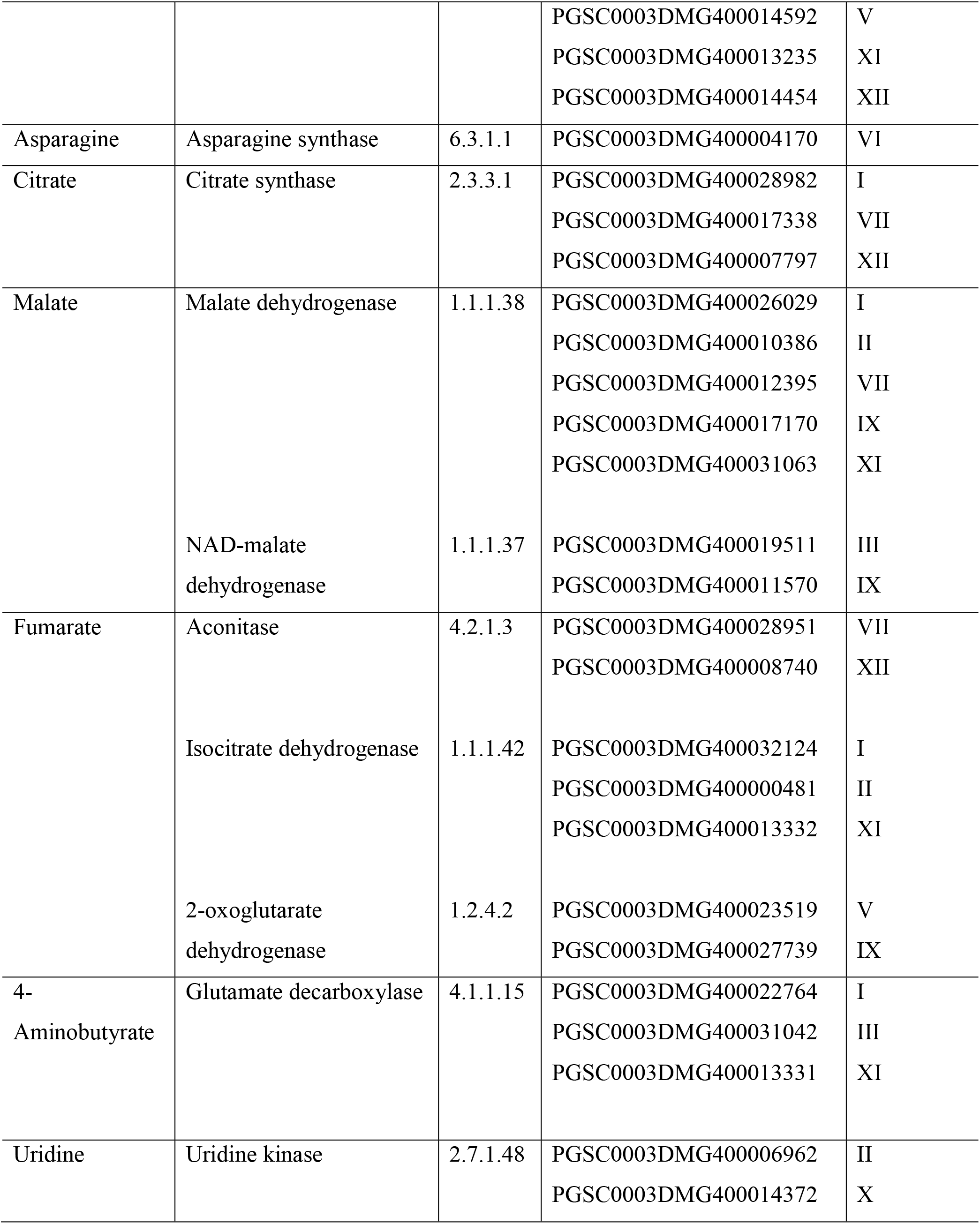
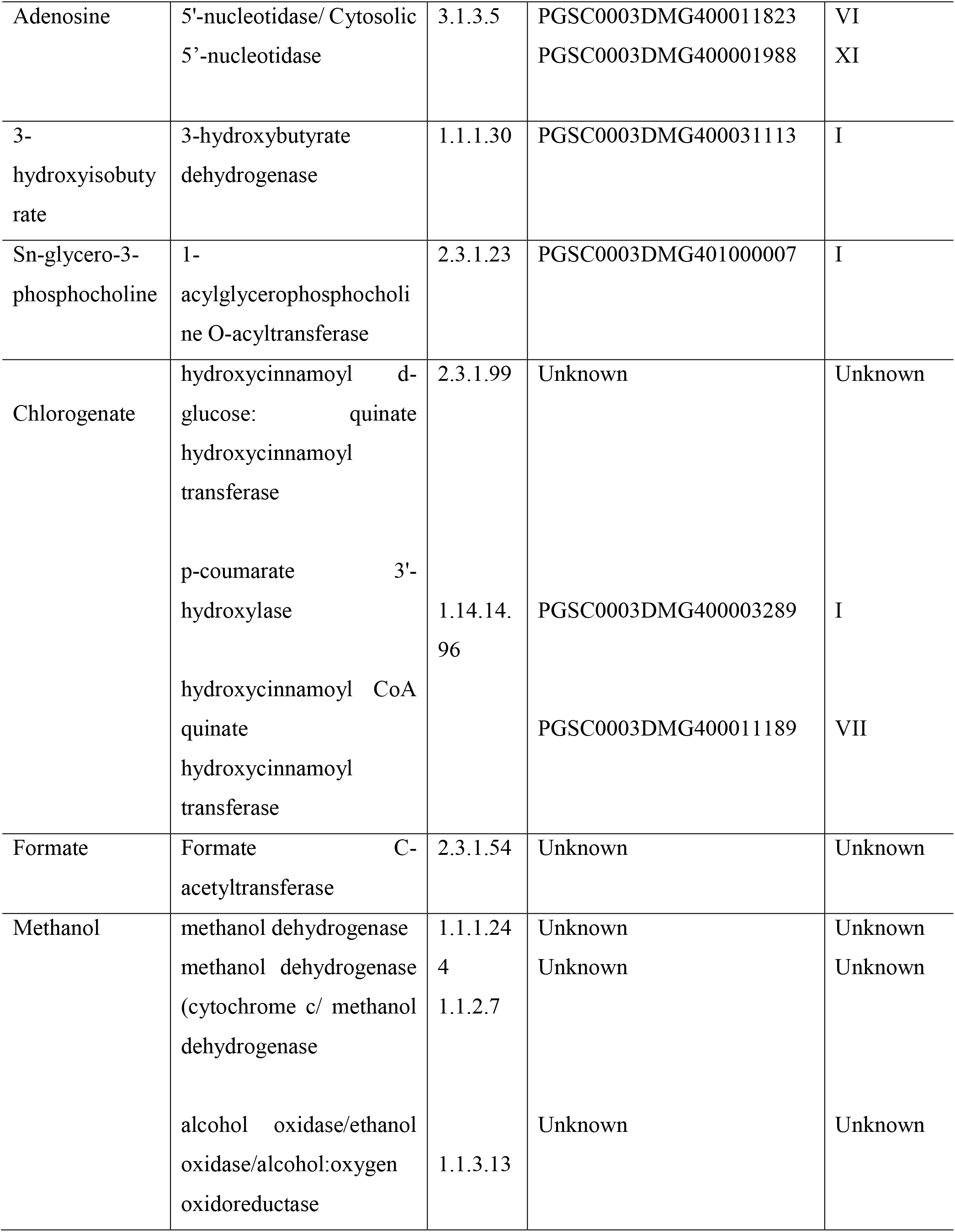
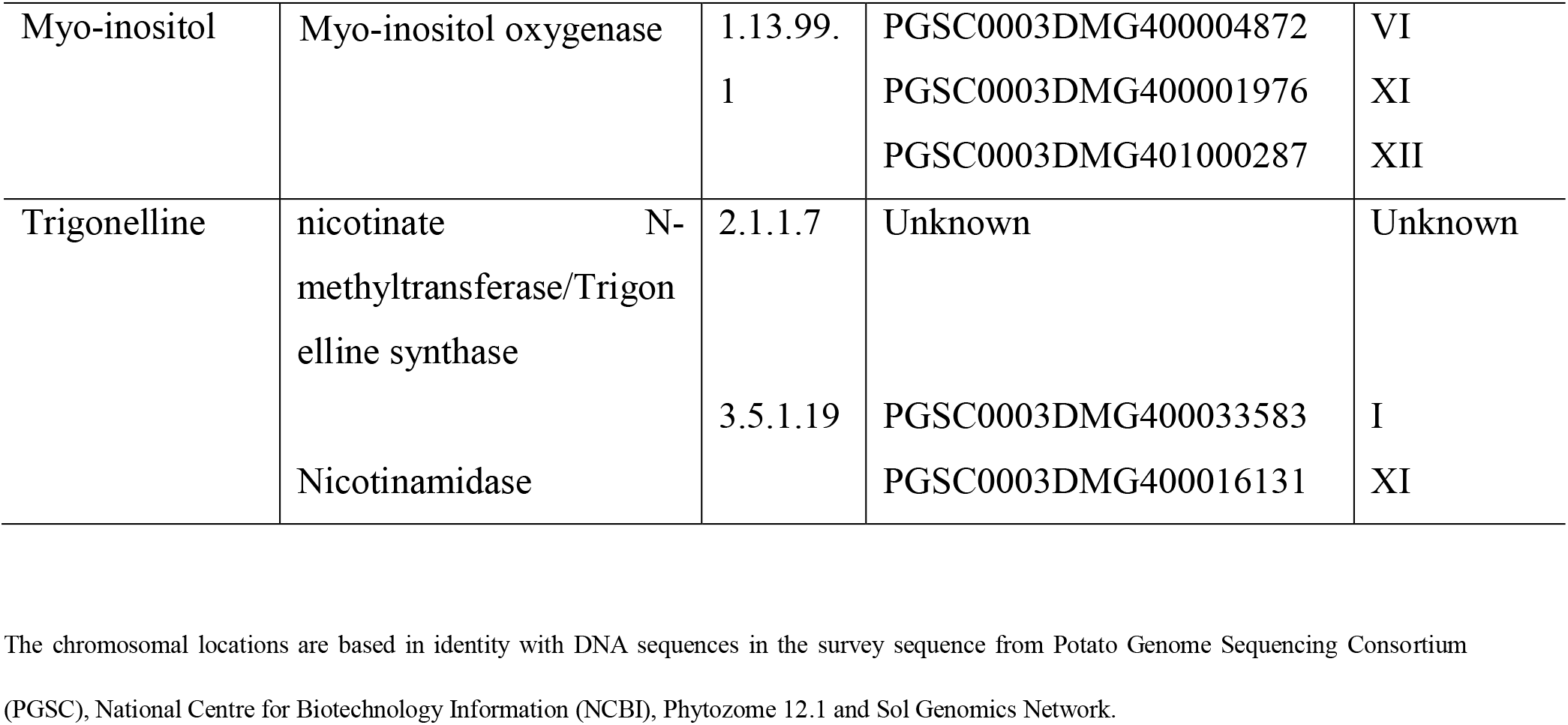
Summary of potato gene sequences with annotation and amino acid identity to enzymes producing significant metabolites identified in the present study.

Considerable variation in the levels of citrate and malate were observed in the present investigation (Fig. 5). Citrate synthase and malate dehydrogenase (Table 1) were mapped on potato chromosomes (Chen *et al.*, 2001). However, the natural allelic variations present in these genes controlling citrate and malate levels have not been documented. It is noticeable that the levels of glutamine were exclusively significantly higher in FL-1533 and Kufri Pukhraj after CS as compared to rest of the potato cultivars (Fig. 5, Fig. 6) probably due to an elevated transcription of glutamine synthetase (Roessner-Tunali *et al.*, 2003). Enzymes branched-chain amino acid aminotransferase and glutamine synthetase (Table 1) involved in glutamine biosynthesis were found to be potentially involved in potato tuber quality traits (Ducreux *et al.*, 2008). Significantly increased asparagine levels were observed in Atlantic, Kufri Jyoti, Kufri Pukhraj, and PU1 after CS (Fig. 3, Fig. 4, and Fig. 5). Silencing of vacuolar invertase and asparagine synthetase (*AS1* and *AS2*) genes demonstrated that the transcript levels of these genes were correlated with RS and asparagine content in transgenic (Zhu *et al.*, 2016). Therefore, metabolite variations (Fig. 3, Fig. 4, and Fig. 5) and various correlations (Supplementary Fig. S3-S7) obtained especially after cold storage raises the possibility to test the function of genes or the combination of transgenes using genetic engineering approaches to further validate the role of other candidate genes identified in this study. Also, the future challenge will be to perform the qRT-PCR assays to ascertain and discover the expression of genes involved in CIS and finding their precise role in controlling metabolite accumulation. This approach will allow the identification of new candidate genes involved in CIS processes and can be used in further genetic improvement of potato tuber quality.

## Conclusions

A number of commercial potato cultivars used for processing and table purpose are currently available. However, the information on physiological, biochemical, and molecular mechanisms underlying the CIS status of various potato cultivars is very scanty. So far much attention has been given towards understanding the role of RS and asparagine in the CIS process and processing attributes of potato tubers. Here, we have presented the differences in natural variation in several other tuber metabolite contents such as amino acids, citrate, malate, methanol, etc. especially after cold storage which indicated that these metabolites can be used to distinguish potato cultivars differing in their CIS response and processing quality attributes. Selection and development of potato cultivars for long term storage along with good processing attributes using traditional breeding techniques may be cumbersome. Hence, the knowledge of appropriate parents using metabolite diversity is needed. Also, the presence or absence of specific metabolites cannot be the only concluding answer for the prediction of CIS behaviour, therefore, the relative amount of the specific metabolite present can also play a contribution in CIS status of potato cultivar. Therefore, the knowledge and information of various metabolites along with candidate genes are necessary for the detailed understanding of various biochemical mechanisms underlying metabolite variations in different potato cultivars differencing in their CIS response. Information obtained based on such observations is of major interest to potato breeders and the processing industry for further utilization of metabolite marker in the selection of CIS resistant potato genotypes.

## Acknowledgements

The research was supported by the Department Research and Development Program (DRDP), Department of Biotechnology, Savitribai Phule Pune University. The authors are also grateful to BT Company; and Jai Kisan Farm Products and Cold Chains Pvt. Ltd, India, Pune for generously providing the potato cultivars. The authors acknowledge HF-NMR facility at IISER-Pune (co-funded by DST-FIST and IISER Pune). SY is thankful for the financial assistance from UGC-JRF, Government of India. JC acknowledges the funding from IISER Pune, Government of India; extramural funding from the Science and Engineering Research Board (SERB), Govt. of India (EMR/2015/001966), and from Department of Biotechnology (DBT), Govt. of India (BT/PR24185/BRB/10/1605/2017). SS acknowledges the funding from Ramalingaswami fellowship (BT/RLF/Re-entry/11/2012; Department of Biotechnology - DBT, Government of India); and University Grants Commission (UGC, Government of India F.4-5(18-FRP) (IV-Cycle)/2017(BSR)). MK acknowledges DBT, GOI for his Masters in Biotechnology fellowship.

## Conflict of interest

Authors declare no potential conflict of interest.

