## Supplementary Tables and Supplementary Figures for "Cold storage reveals distinct metabolic perturbations in processing and non-processing cultivars of potato"

**Title:** Cold-storage reveals distinct metabolite perturbations in processing and non-processing cultivars of potato (*Solanum tuberosum* L.)

**Authors:** Sagar S Datir*[,§,1,a], Saleem Yousf[§,3], Shilpy Sharma[1], Mohit Kochle[1], Ameeta Ravikumar[2], and Jeetender Chugh*[,3,4]

**The affiliations and addresses of the authors:**

[1]Department of Biotechnology, Savitribai Phule Pune University, Pune – 411007, India

[a]Present address: Biology Department, Biosciences Complex, Queen's University, Kingston, ON, CA K7L 3N6

[2]Institute of Bioinformatics and Biotechnology, Savitribai Phule Pune University, Pune – 411007, India

[3]Department of Chemistry, and [4]Department of Biology, Indian Institute of Science, Education and Research, Pune – 411008, India

***Corresponding authors:**

Sagar S Datir, Ph.D.

Address: Department of Biotechnology, Savitribai Phule Pune University, Pune – 411007, India

ORCID: 0000-0003-0065-498X

and

Jeetender Chugh, Ph.D.

Assistant Professor, Department of Chemistry & Biology,

C-115, Indian Institute of Science Education & Research, Dr. Homi Bhabha Road, Pashan, Pune – 411008, India

ORCID: 0000-0002-9996-5202

**Supplementary Table 1**

**Information on potato cultivars**

| Variety / Cultivar | Source | Type | Shape | Colour | | Reducing sugar content (g/100g) | Total sugars (g/100g) | Dry matter content (g/100g) | Starch content (g/100g) | Storage behaviour |
|---|---|---|---|---|---|---|---|---|---|---|
| | | | | Skin | Flesh | | | | | |
| Frito Lay-1533 | Pepsi Foods Pvt. Ltd. Channo, Sangrur | Processing | Oval | Light russet | White | 0.13 | 1.70 | 21.75 | 55.10 | Good |
| Atlantic | Pepsi Foods Pvt. Ltd. Channo, Sangrur | Processing | Round | Brownish yellow | White | 0.35 | 0.63 | 24.32 | 63.12 | Good |
| Kufri Pukhraj | Central Potato Research Institute, Shimla | Non-processing | Oval | Brown | Cream | 0.67 | 3.29 | 16.72 | 58.99 | Average/Poor |
| Kufri Joyti | Central Potato Research Institute, Shimla | Non-processing | Round | Brownish yellow | Cream | 0.87 | 1.98 | 17.47 | 51.52 | Medium/Poor |
| PU1 | No information | -- | Oval | Brownish yellow | Cream | 0.02 | 0.41 | -- | -- | Medium/Poor |

The information is adapted from Aggarwal *et al*., 2017, Kaur and Khurana 2017; Datir *et al*., 2019

**Supplementary Table S2**

**List of the metabolites detected**

| S. No. | Metabolite | CID Code | Chemical shift | Assignment |
|---|---|---|---|---|
| 1 | 3-hydroxyisobutyrate | 87 | 1.07 (d) | β-$CH_3$ |
| | | | 2.47 (m) | α-CH |
| 2 | 4-Aminobutyrate | 119 | 1.89 (m) | β-$CH_2$ |
| | | | 2.29 (t) | α-$CH_2$ |
| | | | 3.00 (t) | γ-$CH_2$ |
| 3 | Adenosine | 60961 | 6.07 (d) | $C_2H$ |
| 4 | Alanine | 5950 | 3.80 (q) | α-CH |
| | | | 1.46 (d) | β-$CH_3$ |
| 5 | Allantoin | 204 | 5.37 (s) | CH |
| 6 | Arginine | 6322 | 1.68 (m) | γ-$CH_2$ |
| | | | 1.89 (m) | β-$CH_2$ |
| | | | 3.23 (t) | δ-$CH_2$ |
| | | | 3.76 (t) | α-CH |
| 7 | Ascorbate | 54670067 | 4.51 (d) | $C_4H$ |
| 8 | Asparagine | 6267 | 2.84 (m) | β-$CH_2$ |
| | | | 2.94 (m) | β-$CH_2$ |
| | | | 3.99 (dd) | α-CH |
| 9 | Aspartate | 5960 | 2.66 (dd) | β-$CH_2$ |
| | | | 2.79 (dd) | β-$CH_2$ |
| | | | 3.91 (dd) | α-CH |
| 10 | Chlorogenate | 1794427 | 6.40 (d) | $C_{15}H$ |
| | | | 6.95 (d) | $C_{21}H$ |
| | | | 7.13 (dd) | $C_{22}H$ |
| | | | 7.19 (d) | $C_{18}H$ |
| | | | 7.66 (d) | $C_{16}H$ |
| 11 | Choline | 305 | 3.20 (s) | $N(CH_3)_3$ |
| | | | 3.52 (m) | $NCH_2$ |
| | | | 4.06 (m) | $OCH_2$ |
| 12 | Citrate | 311 | 2.67 (d) | $CH_2$ |
| | | | 2.76 (d) | $CH_2$ |
| 13 | DSS | 74873 | 0.00 (s) | $Si(CH_3)_3$ |

| | | | | |
|---|---|---|---|---|
| | | | 0.62 (t) | γ-$CH_2$ |
| | | | 1.75 (m) | β-$CH_2$ |
| | | | 2.91 (t) | α-$CH_2$ |
| 14 | Formate | 283 | 8.44 (s) | CH |
| 15 | Fructose | 2723872 | 3.54 (m) | $C_1H$ |
| | | | 3.70 (m) | $C_1H$ |
| | | | 3.88 (dd) | $C_3H$ |
| | | | 4.01 (m) | $C_4H$ |
| | | | 4.11 (d) | $C_5H$ |
| 16 | Fumarate | 5460307 | 6.50 (s) | CH |
| 17 | Galactose | 6036 | 4.57 (d) | $C_2H$ |
| | | | 5.25 (d) | $C_2H$ |
| 18 | Glucose | 5793 | 3.23 (dd) | $C_3H$ |
| | | | 3.39 (m) | $C_5H$ |
| | | | 3.45 (m) | $C_6H$ |
| | | | 3.52 (m) | $C_3H$ |
| | | | 3.72 (m) | $C_4H/C_{11}H$ |
| | | | 3.82 (m) | $C_{11}H/C_6H$ |
| | | | 3.88 (dd) | $C_{11}H$ |
| | | | 4.63 (d) | $C_2H$ |
| | | | 5.22 (d) | $C_2H$ |
| 19 | Glutamate | 33032 | 2.03 (m) | β-$CH_2$ |
| | | | 2.10 (m) | β-$CH_2$ |
| | | | 2.34 (m) | γ-$CH_2$ |
| | | | 3.75 (dd) | α-CH |
| 20 | Glutamine | 5961 | 2.14 (m) | β-$CH_2$ |
| | | | 2.459 (m) | γ-$CH_2$ |
| | | | 3.76 (t) | α-CH |
| 21 | Glycine | 750 | 3.56 (s) | $CH_2$ |
| 22 | Isoleucine | 6306 | 0.92 (t) | δ-CH3 |
| | | | 0.99 (d) | β-$CH_3$ |
| | | | 1.24 (m) | γ-$CH_2$ |
| | | | 1.45 (m) | γ-$CH_2$ |
| | | | 1.97 (m) | β-CH |

| | | | | |
|---|---|---|---|---|
| | | | 3.66 (d) | α-CH |
| 23 | LDL | | 0.84 (t) | $CH_3(CH_2)_n$ |
| | | | 1.25 (m) | $(CH_2)_n$ |
| 24 | Leucine | 6106 | 0.94 (d) | δ-CH3 |
| | | | 0.95 (d) | δ-CH3 |
| | | | 1.68 (m) | β-CH2 |
| | | | 1.70 (m) | γ-CH |
| | | | 3.71 (t) | α-CH |
| 25 | Lysine | 5962 | 1.49 (m) | γ-$CH_2$ |
| | | | 1.72 (m) | δ-$CH_2$ |
| | | | 1.89 (m) | β-$CH_2$ |
| | | | 3.01 (t) | e-$CH_2$ |
| | | | 3.74 (t) | α-CH |
| 26 | Malate | 525 | 2.37 (dd) | β-$CH_2$ |
| | | | 2.66 (dd) | β-$CH_2$ |
| | | | 4.29 (dd) | α-CH |
| 27 | Mannose | 18950 | 4.89 (d) | $C_2H$ |
| | | | 5.17 (d) | $C_2H$ |
| 28 | Methanol | 887 | 3.34 (s) | $CH_3$ |
| 29 | Methionine | 6137 | 2.12 (s) | S-$CH_3$ |
| | | | 2.17 (m) | β-$CH_2$ |
| | | | 2.63 (t) | S-$CH_2$ |
| | | | 3.85 (t) | α-CH |
| 30 | Myo-inositol | 892 | 3.27 (t) | $C_5H$ |
| | | | 3.54 (dd) | $C_1H/C_3H$ |
| | | | 3.62 (dd) | $C_4H/C_6H$ |
| | | | 4.05 (t) | $C_2H$ |
| 31 | Phenylalanine | 6140 | 7.42 (m) | $C_3H/C_5H$ |
| | | | 7.36 (m) | $C_2H/C_6H$ |
| | | | 7.32 (m) | $C_4H$ |
| | | | 3.98 (dd) | α-CH |
| | | | 3.27 (m) | β-$CH_2$ |
| 32 | Proline | 145742 | 4.11 (t) | α-CH |
| | | | 3.40 (m) | δ-$CH_2$ |
| | | | 3.33 (m) | δ-$CH_2$ |

| | | | | |
|---|---|---|---|---|
| | | | 2.34 (m) | β-$CH_2$ |
| | | | 2.07 (m) | β-$CH_2$ |
| | | | 2.01 (m) | γ-$CH_2$ |
| 33 | Pyroglutamate | 7405 | 2.02 (m) | $C_4H$ |
| | | | 2.40 (m) | $C_3H$ |
| | | | 2.49 (m) | $C_4H$ |
| | | | 4.16 (m) | $C_5H$ |
| 34 | Serine | 5951 | 3.83 (m) | α-CH |
| | | | 3.94 (m) | β-$CH_2$ |
| | | | 3.98 (m) | β-$CH_2$ |
| 35 | sn-glycero-3-phosphocholine | 657272 | 3.21 (s) | $C_{13}H/C_{14}H$ |
| | | | 4.31 (m) | $C_3H$ |
| 36 | Sucrose | 5988 | 3.46 (t) | $C_{10}H$ |
| | | | 3.55 (m) | $C_{12}H$ |
| | | | 3.75 (m) | $C_{11}H$ |
| | | | 3.81 (m) | $C_{17}H/C_{19}H$ |
| | | | 3.88 (m) | $C_5H$ |
| | | | 4.04 (t) | $C_4H$ |
| | | | 4.20 (d) | $C_3H$ |
| | | | 5.40 (d) | $C_7H$ |
| 37 | Threonine | 6288 | 1.32 (d) | γ-$CH_3$ |
| | | | 3.58 (d) | α-CH |
| | | | 4.24 (m) | β-$CH_2$ |
| 38 | Trigonelline | 5570 | 4.42 (s) | $C_9H$ |
| | | | 8.07 (t) | $C_4H$ |
| | | | 8.82 (m) | $C_5H/C_3H$ |
| | | | 9.11 (s) | $C_1H$ |
| 39 | Trypotophan | 6305 | 3.29 (m) | $CH_2$ |
| | | | 4.05 (m) | CH |
| | | | 7.19 (t) | $C_5H/C_6H$ |
| | | | 7.27 (t) | $C_5H/C_6H$ |
| | | | 7.31 (s) | $C_2H$ |
| | | | 7.53 (d) | $C_7H$ |

| | | | | |
|---|---|---|---|---|
| | | | 7.72 (d) | $C_4H$ |
| 40 | Tyrosine | 6057 | 7.17 (d) | $C_2H/C_6H$ |
| | | | 6.87 (d) | $C_3H/C_5H$ |
| | | | 3.93 (dd) | α-CH |
| | | | 3.18 (dd) | β-$CH_2$ |
| | | | 3.04 (dd) | β-$CH_2$ |
| 41 | Uridine | 6029 | 7.85 (d) | $C_{11}H$ |
| | | | 5.9 (d) | $C_2H$ |
| | | | 5.89 (d) | $C_{10}H$ |
| | | | 4.34 (dd) | $C_3H$ |
| | | | 4.22 (dd) | $C_4H$ |
| | | | 4.12 (m) | $C_5H$ |
| | | | 3.9 (dd) | $C_{14}H$ |
| | | | 3.8 (dd) | $C_{14}H$ |
| 42 | Valine | 6287 | 0.99 (d) | $CH_3$ |
| | | | 1.04 (d) | $CH_3$ |
| | | | 2.28 (m) | β-CH |
| | | | 3.61 (d) | α-CH |
| 43 | U1 | | 8.22 (m) | |
| | | | 9.00 (d) | |
| | | | 9.10 (d) | |
| | | | 9.33 (s) | |
| 44 | U2 | | 7.27 (s) | |
| | | | 8.39 (s) | |
| 45 | U3 | | 3.85 | |
| | | | 3.95 | |
| | | | 4.02 | |
| | | | 5.14 (d) | |
| 46 | U4 | | 3.82 | |
| | | | 3.89 | |
| | | | 3.97 | |
| | | | 4.03 | |
| | | | 4.99 | |
| 47 | U5 | | 3.66 | |
| | | | 4.29 (dd) | |

**Supplementary Table S3:** Number of correlation observed for the metabolites in the different cultivars used in the study at two time points – fresh harvest and one month cold storage.

| **Cultivar – time point** | **Positive** | **Negative** | **Total** |
|---|---|---|---|
| Atlantic FH | 40 | 9 | 49 |
| Atlantic CS | 51 | 4 | 55 |
| Frito Lay 1533 FH | 46 | 5 | 51 |
| Frito Lay 1533 CS | 44 | 9 | 53 |
| Kufri Pukhraj FH | 31 | 5 | 36 |
| Kufri Pukhraj CS | 37 | 15 | 52 |
| Kufri Jyoti FH | 43 | 5 | 48 |
| Kufri Jyoti CS | 38 | 15 | 53 |
| PU1 FH | 42 | 3 | 45 |
| PU1 CS | 18 | 13 | 31 |

**Supplementary Table S4:** Metabolite biomarker for cold-induced sweetening.

| **Biomarker name** | **KEGG ID** |
| --- | --- |
| Glucose | C00031 |
| Fructose | C00095 |
| Asparagine | C00152 |
| Citrate | C00158 |
| Malate | C00149 |
| Methanol | C00132 |
| Total amino acids | -- |
| Ratio of asparagine to reducing sugars | -- |

### Supplementary Figures

**Figure S1:** $^{1}$H-$^{1}$H TOCSY correlation spectrum of the methanolic extract of fresh harvest of Kufri Pukhraj potato cultivar (cold storage) used in the study (as described in Materials and Methods). The cross-peaks in the TOCSY spectrum have been used to re-confirm the resonance assignments enlisted in Table S2.

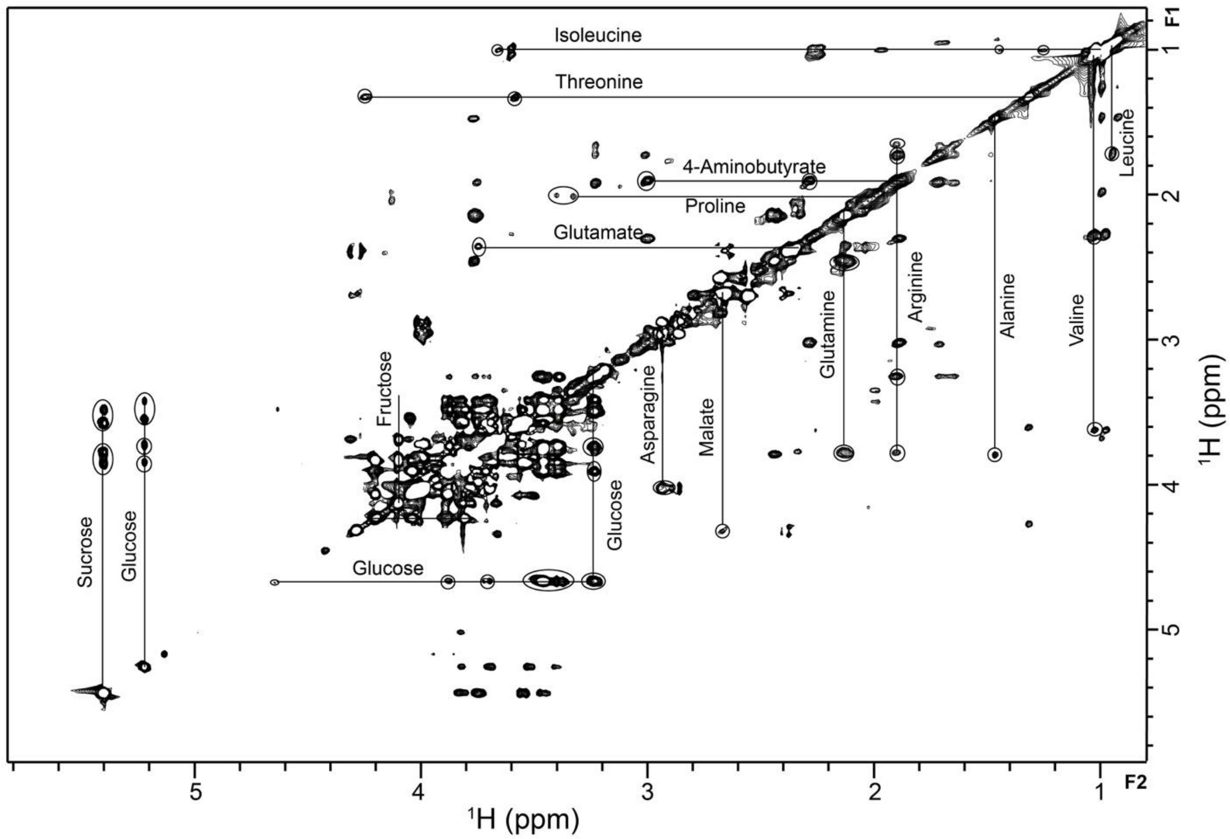

**Figure S2:** 1D $^{1}$H-NMR spectrum of the methanolic extract of the Kufri Pukhraj potato cultivar (Cold storage) used in the study (as described in Materials and Methods). A combination of line-shape, multiplicity, scalar coupling, and chemical shift values obtained from this spectrum were used to identify 39 abundant metabolites. The assignment was validated with the BMRB and HMDB databases. The chemical shift and multiplicity details of the metabolic ensemble (marked in numbers) in the spectrum have been listed in Table S2.

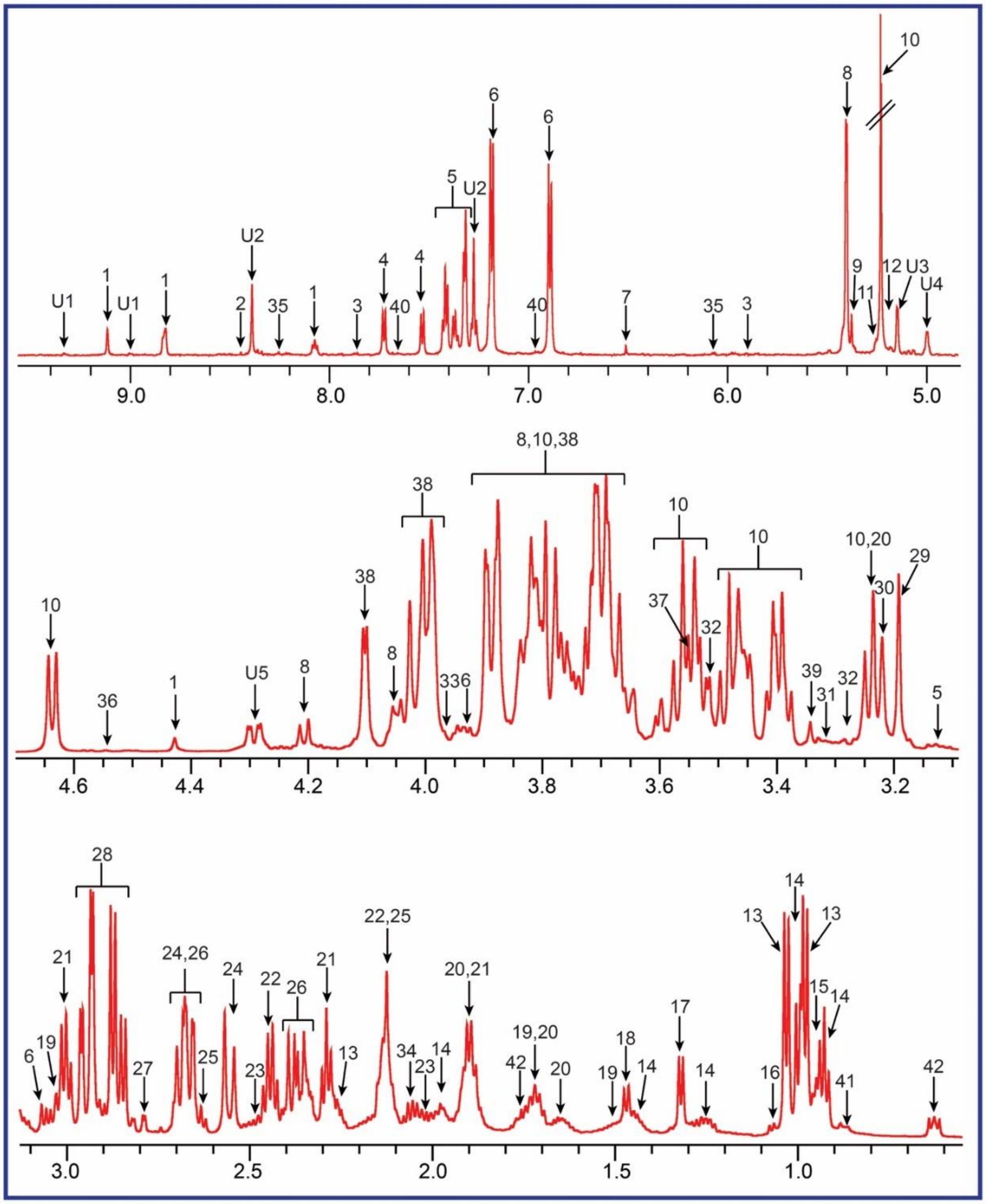

**Figure S3:** Correlation plots between fresh harvest (FH; upper-right half of the plot marked in white) and cold storage (CS; lower-left half of the plot marked in light blue) metabolites for potato tubers of Atlantic cultivar

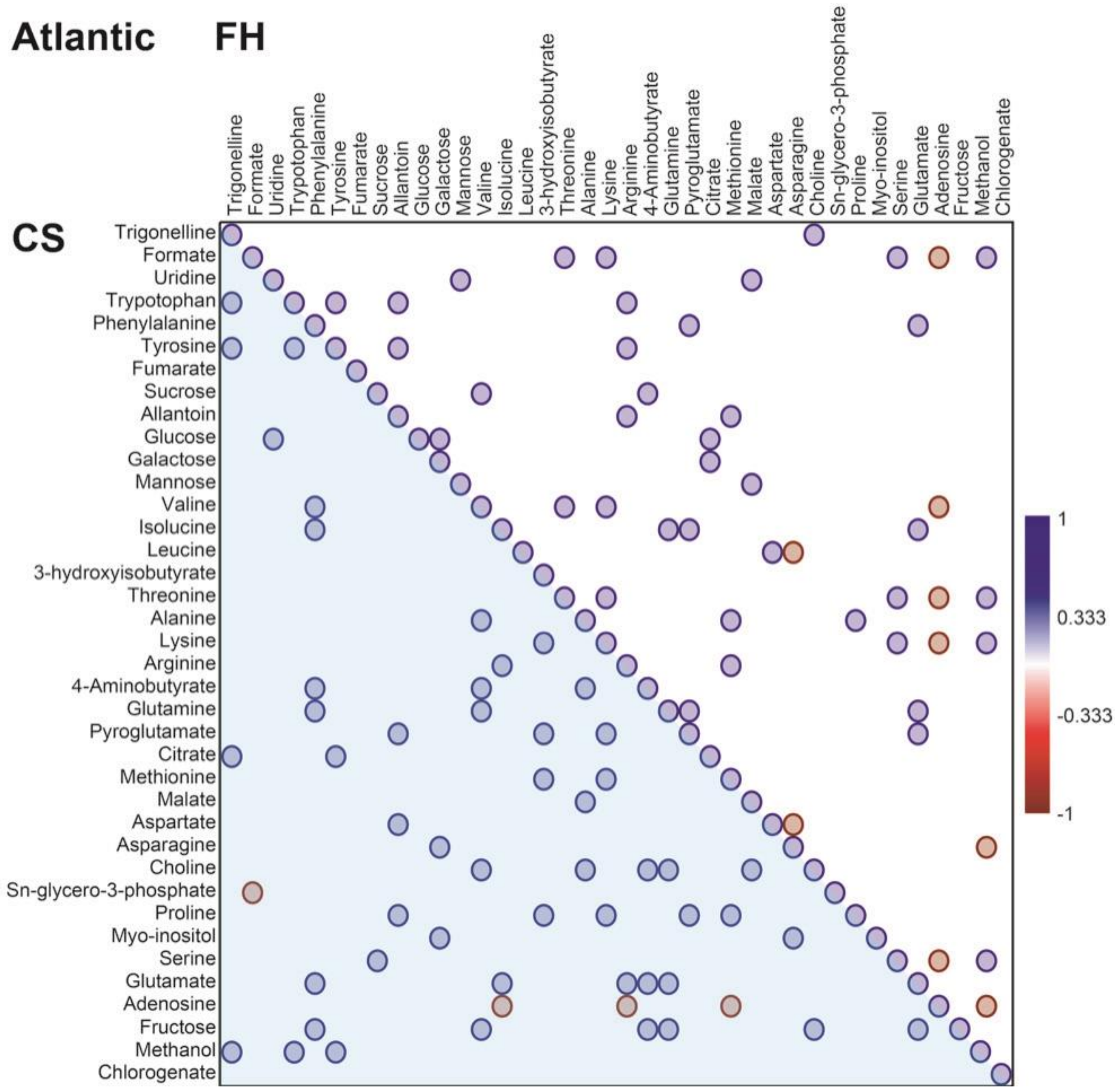

**Figure S4:** Correlation plots between fresh harvest (FH; upper-right half of the plot marked in white) and cold storage (CS; lower-left half of the plot marked in light blue) metabolites for potato tubers of FL-1533 cultivar

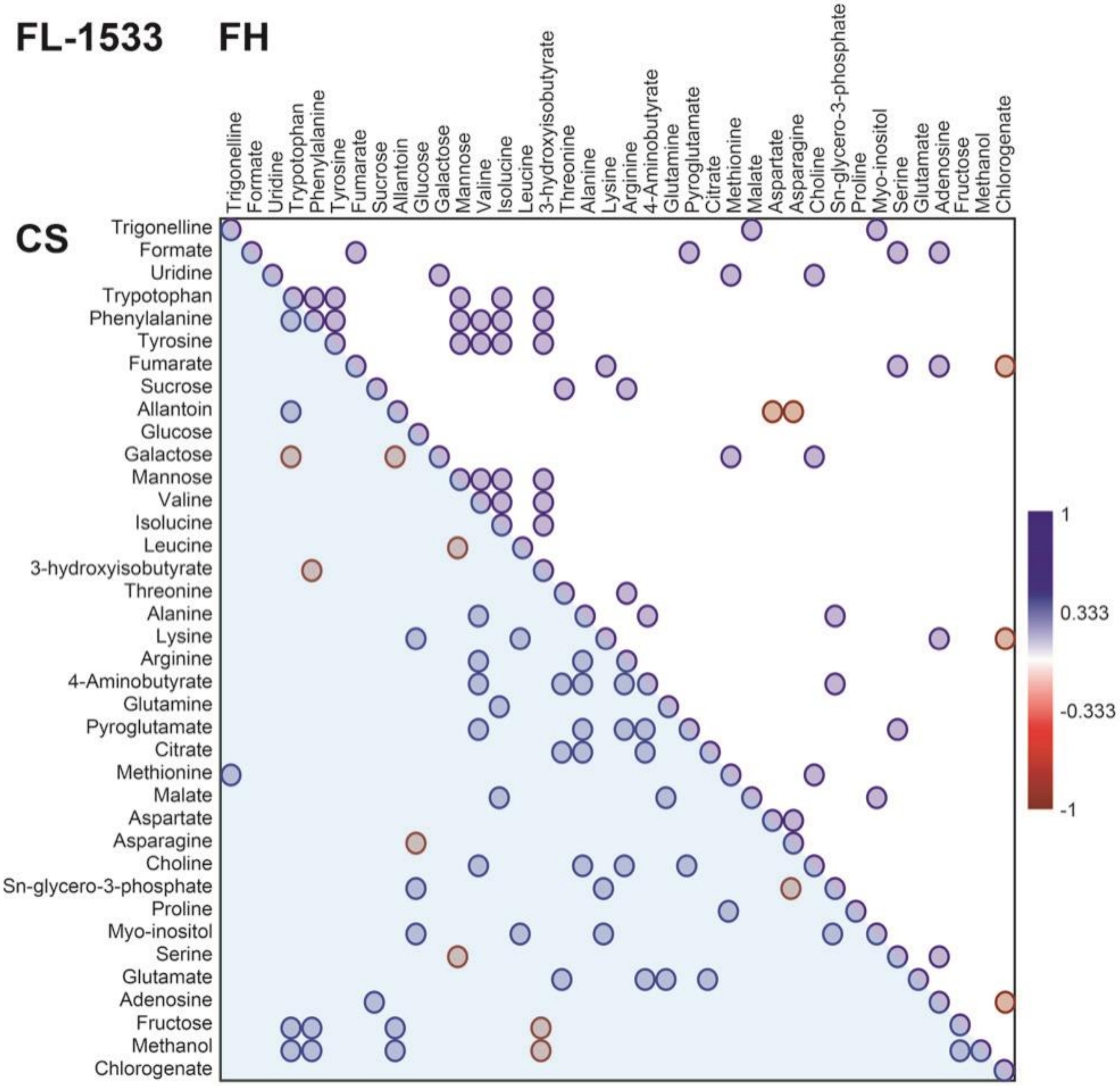

**Figure S5:** Correlation plots between fresh harvest (FH; upper-right half of the plot marked in white) and cold storage (CS; lower-left half of the plot marked in light blue) metabolites for potato tubers of Kufri Jyoti cultivar

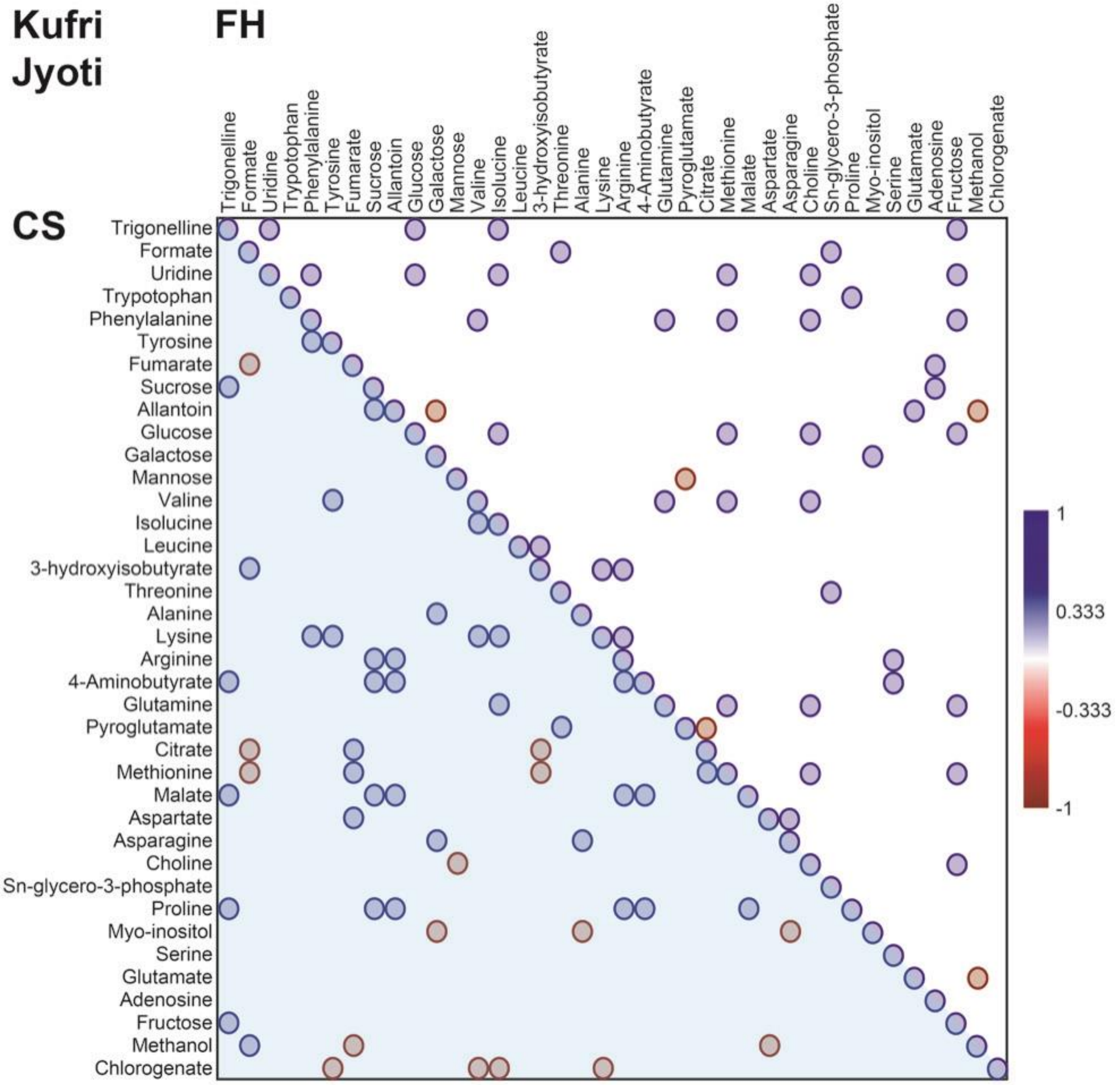

**Figure S6:** Correlation plots between fresh harvest (FH; upper-right half of the plot marked in white) and cold storage (CS; lower-left half of the plot marked in light blue) metabolites for potato tubers of Kufri Pukhraj cultivar

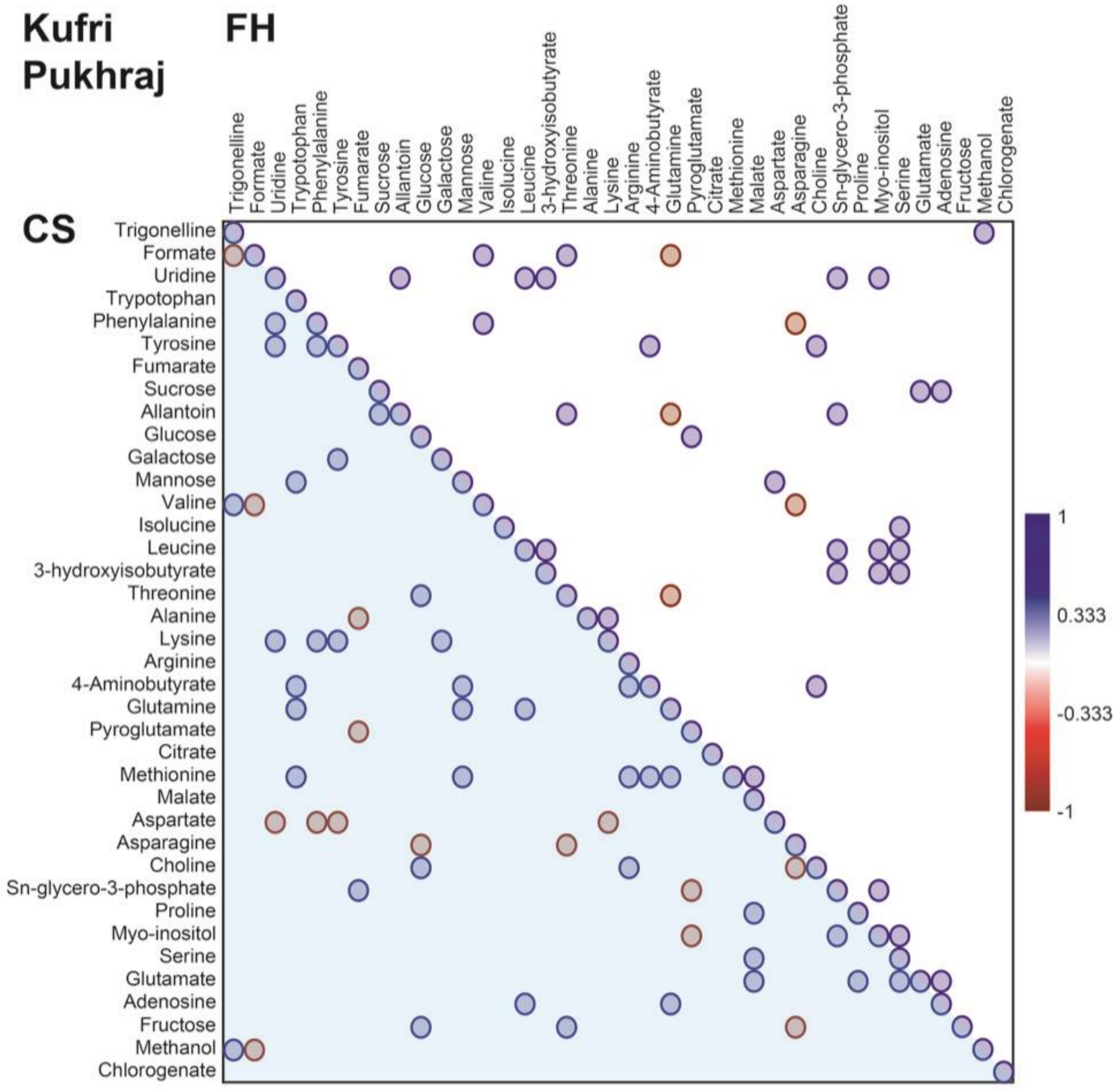

**Figure S7:** Correlation plots between fresh harvest (FH; upper-right half of the plot marked in white) and cold storage (CS; lower-left half of the plot marked in light blue) metabolites for potato tubers of PU1 cultivar

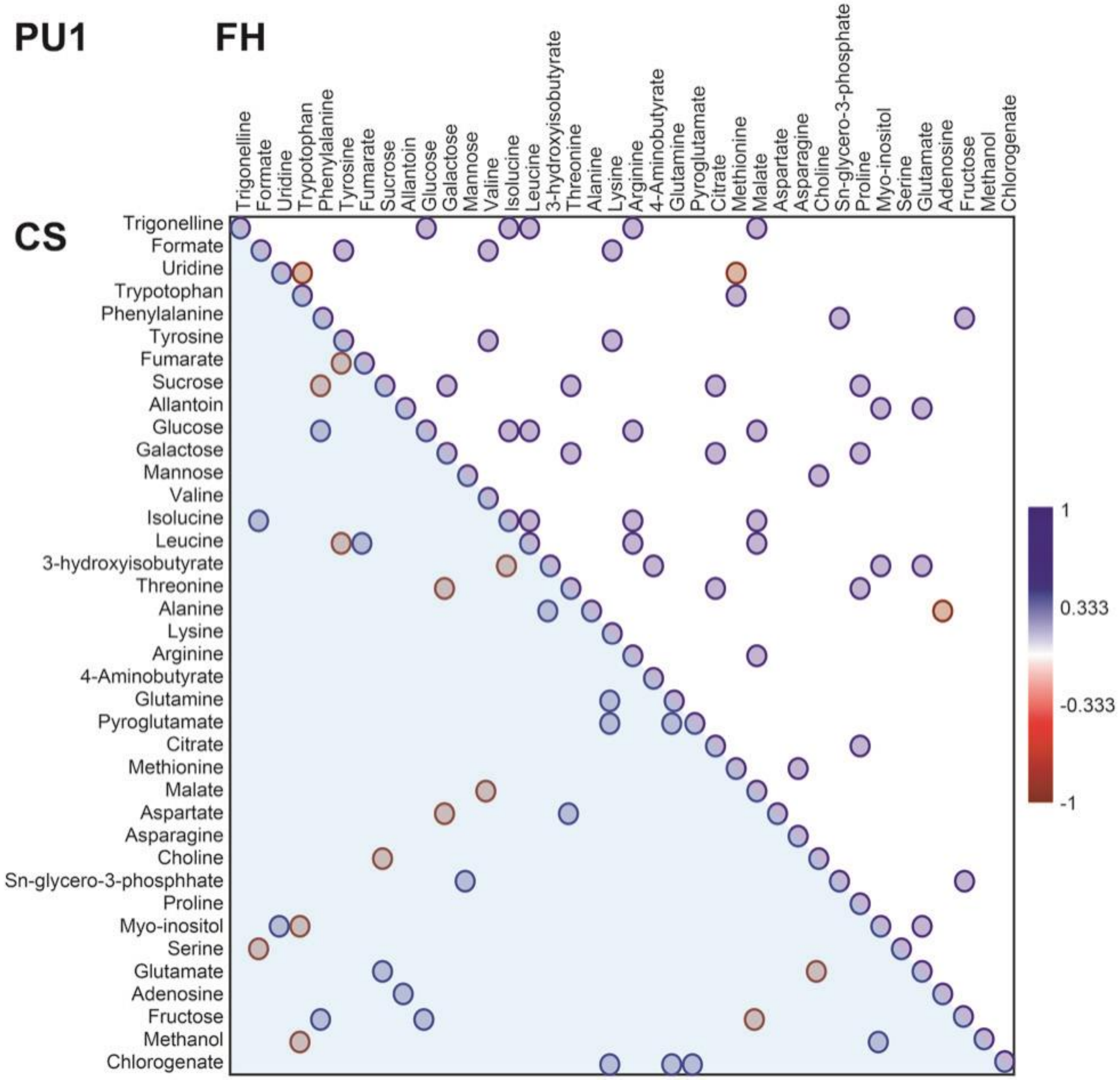
